# The Crohn’s disease-associated *Escherichia coli* strain LF82 rely on SOS and stringent responses to survive, multiply and tolerate antibiotics within macrophages

**DOI:** 10.1101/551226

**Authors:** Gaëlle Demarre, Victoria Prudent, Hanna Schenk, Emilie Rousseau, Marie-Agnes Bringer, Nicolas Barnich, Guy Tran Van Nhieu, Sylvie Rimsky, Silvia De Monte, Olivier Espéli

**Affiliations:** CIRB – Collège de France, CNRS-UMR724, INSERM U1050, PSL Research University, 11 place Marcelin Berthelot 75005 Paris, France.; Inovarion, Paris, France; Department of Evolutionary Theory, Max Planck Institute for Evolutionary Biology, Plön, Germany; Centre des Sciences du Goût et de l'Alimentation, AgroSup Dijon, CNRS, INRA, Université Bourgogne Franche-Comté, F-21000 Dijon, France; Microbes, Intestin, Inflammation et Susceptibilité de l'Hôte. UMR Inserm/ Université d'Auvergne U1071, USC INRA 2018; Institut de Biologie de l’Ecole Normale Supérieure, Département de Biologie, Ecole Normale Supérieure, CNRS, INSERM, PSL Research University, Paris, France

## Abstract

Adherent Invasive *Escherichia coli* (AIEC) strains recovered from Crohn's disease lesions survive and multiply within macrophages. A reference strain for this pathovar, AIEC LF82, forms microcolonies within phagolysosomes, an environment that prevents commensal *E. coli* multiplication. Little is known about the LF82 intracellular growth status, and signals leading to macrophage intra-vacuolar multiplication. We used single-cell analysis, genetic dissection and mathematical models to monitor the growth status and cell cycle regulation of intracellular LF82. We found that within macrophages, bacteria may replicate or undergo non-growing phenotypic switches. This switch results from stringent response firing immediately after uptake by macrophages or at later stages, following genotoxic damage and SOS induction during intracellular replication. Importantly, non-growers resist treatment with various antibiotics. Thus, intracellular challenges induce AIEC LF82 phenotypic heterogeneity and non-growing bacteria that could provide a reservoir for antibiotic-tolerant bacteria responsible for relapsing infections.

## Introduction

Adherent Invasive *Escherichia coli* (AIEC) strains recovered from Crohn’s disease (CD) lesions are able to adhere to and invade cultured intestinal epithelial cells and to survive and multiply within macrophages (Darfeuille-Michaud *et al*, 1998; Glasser *et al*, 2001). Attention around the potential role of AIEC in the pathophysiology of CD is growing (Elhenawy *et al*, 2018); however much remains to be learned about the host-pathogen interactions that govern AIEC infection biology. The diversity of virulence factors displayed by multiple AIEC strains suggests that members of this pathovar have evolved different strategies to colonize their hosts (Tawfik *et al*, 2014). AIEC ability to persist, and in some cases replicate within macrophages is particularly intriguing. Previous work performed with murine macrophage cell lines has revealed that the prototype AIEC strain LF82, multiplies in a vacuole presenting the characteristics of a mature phagolysosome (Bringer *et al*, 2006; Lapaquette *et al*, 2012). In such an environment, AIEC should encounter acidic, oxidative, genotoxic and proteic stresses. Screening of genes involved in LF82 fitness within macrophage has revealed that HtrA, DsbA, or Fis proteins are required for optimum fitness, (Bringer *et al.*, 2005; Bringer *et al.*, 2007; Miquel *et al.*, 2010). These observations confirmed that LF82 encounter stresses in the phagolysosomes. The impact of these stresses on the survival and growth of LF82 inside phagolysomes has not yet been investigated.

Studies on the bacterial cell cycle of few model organisms under well-controlled laboratory conditions have revealed that to achieve accurate transmission of the genetic information and optimal growth of the population, molecular processes must be coordinated. (for reviews see Hajduk *et al.*, 2016; Haeusser & Levin, 2008). When growth conditions deteriorate, the cell cycle can be modified slightly, as in the case of cell filamentation when genotoxic stress induces the SOS response, or more drastically when sporulation is induced by nutrient deprivation (Jonas, 2014). Such cell cycle alterations affect the entire population. However, under unperturbed conditions, a subset of the population also appears to present a significantly reduced growth rate that allows tolerance to antibiotic treatments. This small portion of the population, typically 1/10000 bacteria, is known as persisters (Wood *et al.*, 2013; Lewis, 2010; Bigger, 1944). Persisters have been detected for a number of bacteria. They can be found spontaneously in normally growing or stationary phase populations, or they are induced by exogeneous stresses or mutations. Significant increase of the proportion of *S. typhimurium* persisters has been observed when these bacteria invade macrophages (Helaine *et al.*, 2014). Using a fluorescent reporter, it has been demonstrated that these persisters were not multiplying prior to antibiotic addition. Recently, the same tool also revealed the presence of non-growing mycobacteria inside macrophages (Mouton *et al.*, 2016). Several mediators of persistence have been identified, with toxin-antitoxin modules emerging as key players and the reduction of metabolic activities as the main driver of persistence (Rycroft *et al.*, 2018; Dörr *et al.*, 2009; Shan *et al.*, 2017; Balaban *et al.*, 2004; Harms *et al.*, 2017; Amato *et al.*, 2014). Persisters are increasingly viewed as a major cause of the recurrence of chronic infectious disease and could be an important factor in the emergence of antibiotic resistance (Verstraeten *et al.*, 2016). In addition to persisters, bacterial tolerance to antibiotic treatments has been observed. In contrast to persisters, tolerance concerns the entire population. Tolerance corresponds to a weaker ability of antibiotics to kill slow growing compared to fast growing bacteria. Tolerant bacteria emerge, for example, in the presence of a nutritional limitation. The viability of tolerant bacteria is impacted by the concentration and length of the antibiotic challenge (Kim & Wood, 2017). Tolerant cells have some aspect of active metabolism, and their frequency in the population changes when bacterial environmental sensing is altered (Amato & Brynildsen, 2015; Bernier *et al.*, 2013; Radzikowski *et al.*, 2016). Persistence is often viewed as the result of a phenotypic switch ensuring long-term adaptation to variable environments, however the origin of persistence and tolerance *in vivo* remain unclear, and their distinction in the context of a host- pathogen interaction is difficult (Kim & Wood, 2017).

In the present work, we analyzed growth characteristics of the prototype AIEC strain LF82 in THP1 monocyte–derived macrophages. We observed that stresses within macrophages induce a profound bacterial response that leads to the formation of non-growing and antibiotic-tolerant LF82 bacteria at a high rate through the successive induction of stringent and SOS responses. A portion of non-growing LF82 produced within macrophages is tolerant to antibiotics and presents a survival advantage. Our work revealed that internalization within phagolysosomes curbs bacterial multiplication, and frequent escape from the replicative cycle toward non-growing state(s) is a way to improve long-term survival in the host.

## Results

### Inside macrophages, LF82 population size increases despite extensive death

We used THP1 monocyte-derived into macrophages to monitor the population size of LF82 bacteria over a 24 h period post infection (P.I.) (Figure 1A). Colony-forming units (CFU) measurements revealed that the LF82 population exponentially increased for 10-14 hours (τ= 0.15 h^−1^, 0.21 doubling / h) after a long lag. The population reached a maximum at 18-20 h of approximately 5-fold the value at 1 h; the number of LF82 then slightly decreased (reaching 3 -fold) at 24 h. In this environment LF82 might simultaneously encounter acidic pH, oxidative and genotoxic stresses, toxic molecules such as cathepsins and a lack of important nutrients. Surprisingly, the tolerance level of LF82 to any individual stress did not differ from a K12-C600 *E. coli* in *in vitro* conditions (Supplementary Figure S1A). Using direct ex vivo Live and Dead labeling, it has been previously proposed that 80% of LF82 present at 24h within macrophages were alive (Lapaquette *et al.*, 2012). We observed that this method slightly underestimates dead bacteria inside macrophages because of a weak propidium iodide (PI) labeling (Supplementary Figure S2A). Live and dead assay performed immediately after macrophage lysis revealed a nearly constant proportion of dead LF82 in the population (20 - 30%) at 1 h, 12 h, 18h and 24 h post-infection (Figure 1A). To estimate the speed of dead bacteria disappearing in macrophages, we observed the elimination of heat-killed bacteria by THP1 macrophages. Dead LF82 disappeared exponentially with a decay rate of 0.6 h^−1^ and a half-life of 1.4 h (Supplementary Figure S2B); therefore dead LF82 observed at 12h, 18h or 24 h did not correspond to the accumulation over infection period but rather to the bacteria killed in the last 3 hours before observations. This finding led to consider that LF82 must be under stress attack by macrophages at all times during infection.

**Figure 1.**
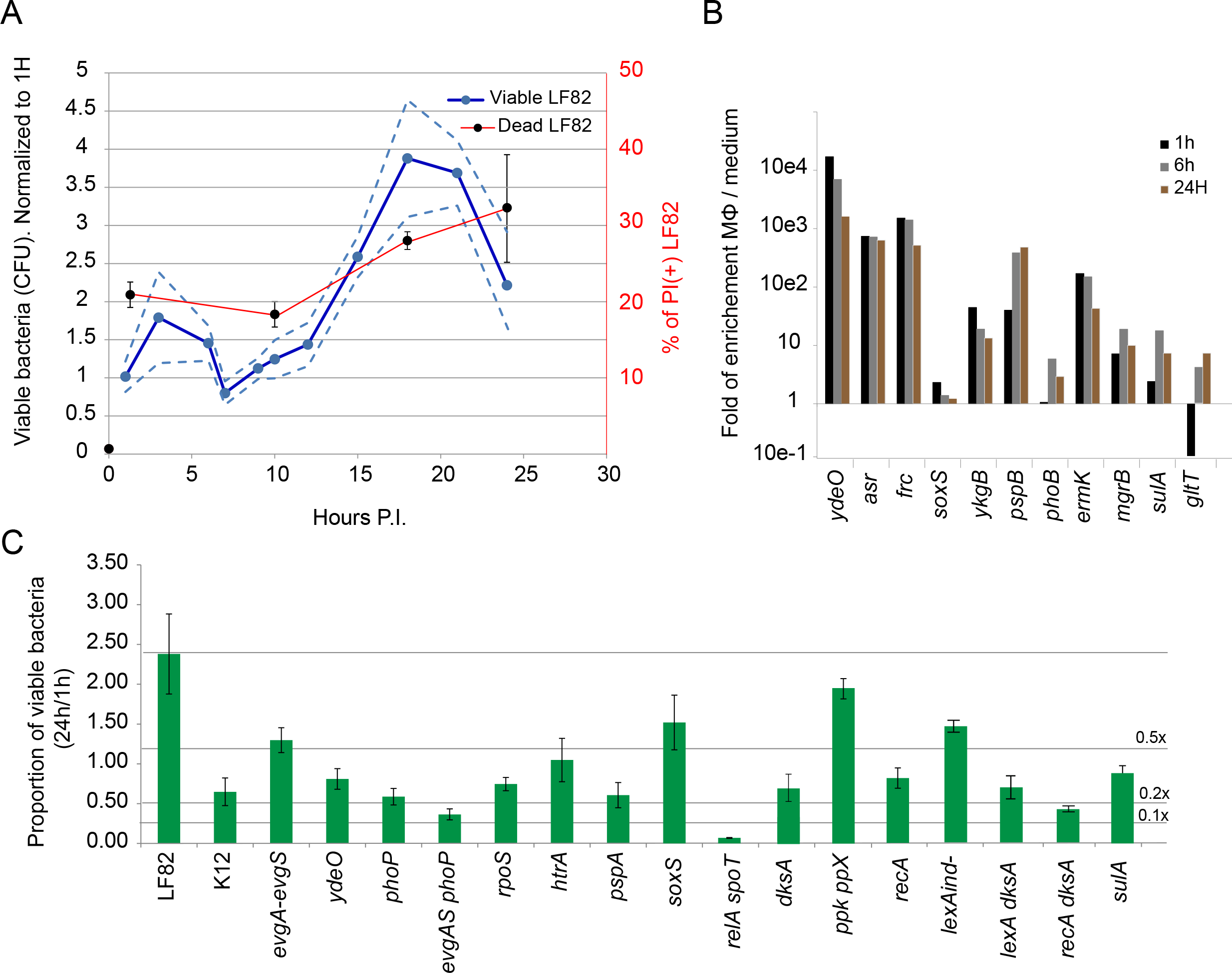
Intracellular LF82 multiplication requires stress responses. A) Measure of viable and dead AIEC LF82 for 24 h post-infection of THP1 differentiated macrophages. Circles represent average CFU (blue) and Propidium iodide (PI) positive bacteria (black) ± Standard deviation (SD) (dotted lines). To minimize variability this experiment has been performed on one set of THP1 amplified for 8 days before differentiation. B) Analysis by qRT-PCR of the induction of LF82 stress responses at 1 h, 6 h and 24 h P.I. of THP1 macrophages. Values represent the average of two experiments (3 technical replicate each). C) Proportion of viable bacteria at 24 h P.I. of THP1 macrophages in comparison to 1 h. LF82, K12-C600 and LF82 deletion mutants were infected at a MOI of 30 which corresponds to 0 - 5 visible bacteria per macrophage at time point 1h (Figure 2A). Values represent the average of 3 to 7 experiments ± SD. Horizontal lines indicate viability decrease by 2, 5 and 10 fold compared to WT LF82.

### LF82 is under attack by macrophages

Using RT-qPCR, we measured the expression of genes induced by the acid (*asr*, *ydeO* and *frc*), oxidative (*soxS* and *ykgB*), and SOS (*sulA*) responses, and the responses to membrane alteration (*pspB*), the lack of Mg2+ (*mgrB*), the lack of phosphate (*phoB*), general efflux pump (*emrK*) and the *gltT* tRNA gene that is repressed by the stringent response (Figure 1B). Every response pathway was induced inside the macrophage. The induction of acid and the oxidative responses was already high at 1 h P.I., while the SOS response, the response to membrane alterations and to the lack of Mg2+, were peaking at 6 h P.I.. The expression of the *gltT* tRNA was strongly repressed at 1 h P.I. indicating that stringent response is on early in the infection. The induction of most pathways decreased at 24 h.

### Environmental stresses influence LF82 survival

To test the impact of stress responses on the ability of LF82 to colonize macrophage, we constructed deletion mutants of several key regulators of *E. coli* stress pathways and analyzed their survival. Deletion of the acid stress regulators *evgA-evgS*, *phoP* and *ydeO* significantly impacted the ability of LF82 to survive and multiply within macrophages to a level comparable to or even below that of a K12-C600 *E. coli* (Figure 1C). Similar observations were obtained with the *rpoS* (general stress response), *recA* (SOS response), *soxS* (oxidative stress) and *pspA* and *htrA* (envelope damages) deletion mutants. The ppGpp0 strain, *relA spoT* deletions, impaired in the stringent response reporting a lack of nutrients, is the most impacted strain; less than 5% of the initial population survived a 24 h period within macrophages. These observations confirmed that LF82 encounter severe stresses in the macrophage environment and that its ability to repair stress mediated injuries will determine survival.

### SOS and stringent responses severely impacted LF82 survival

Because of their known potential to influence growth and cell cycle parameters we explored the stringent and SOS responses in more details. RecA is the main inducer of the SOS response, which activates nearly 100 genes involved in DNA repair and many others with unrelated or unknown functions, but it is also a crucial to correct DNA lesions by homologous recombination and translesion synthesis (Kreuzer, 2013). In addition to *recA* deletion, we constructed deletions of *sulA* (division inhibitor) and a mutation in *lexA* (*lexAind-)*, which blocks SOS induction in MG1655 *E. coli* and reduces viability in the presence of mitomycin C for LF82 and MG1655 (Supplementary Figure S3A). We observed that the deletion of each SOS gene significantly decreased the survival of LF82 within macrophages (Figure 1C). Inside the macrophage, survival of the ppGpp0 strain was dramatically impacted. However, this mutant also presented a strong growth defect in liquid culture that complicates interpretation of the macrophage results. To study the impact of the stringent response on LF82 survival and induced antibiotic tolerance, we constructed deletion mutants that might have partial stringent response phenotypes; deletion of *dksA*, encoding a protein linking the stringent response to transcription (Sharma & Chatterji, 2010); and deletion of the polyphosphate kinase and exopolyphosphatase *ppk* and *ppx* (Rao & Kornberg, 1999). As expected, the *dksA and ppk-ppx* deletions had a much less dramatic effect on LF82 growth and survival within macrophages than the *relA-spoT* mutant; nevertheless, the *dksA* mutation significantly impacted the number of live bacteria recovered at 24 h P.I. (Figure 1C). We investigated the ability of LF82 to survive within macrophages when both stringent and SOS responses were altered. We chose to combine *dksA* deletion with *recA* deletion or *lexAind-* mutation. These strains presented a survival defect comparable to that of the single *dksA* mutant (Figure 1C). These observations demonstrate that surviving LF82 simultaneously or successively require SOS and stringent responses.

### Inside phagolysosomes individual LF82 were not homogenously responding to stresses

Imaging revealed great heterogeneity in the number of LF82 bacteria within individual macrophages. At 18 h or 24 h P.I., many macrophages presented fewer than 5 bacteria, which was comparable to the amount observed at 1 h P.I. (Figure 2A); however, a number of macrophages also presented foci containing up to 50 bacteria (Figure 2B). These observations led us to consider that LF82 were not homogeneously stressed by macrophages. We used GFP fusion with selected stress response promoters to monitor variability of these responses among bacteria and among macrophages (Figure 2B). For all stress responses, heterogeneity in GFP fluorescence for intracellular bacteria was far larger than for LB cultures (Figure 2 F - G). At 24 h P.I., we found that approximately 30% of the bacteria had poorly responded to oxidative or acid responses (Figure 2E and 2F). Owing the high stability of the GFP protein that we used in this assay, it is unlikely that this heterogeneity resulted from short pulses of induction separated by long repression periods. Such a sizeable phenotypic heterogeneity was moreover observed for bacteria contained in a single macrophage as well as similar Lamp 1 positive vacuolar environments (Figure 2C). We therefore questioned whether heterogeneity in stress response might reflect the coexistence of multiple cell cycle regulation phenotypes.

**Figure 2.**
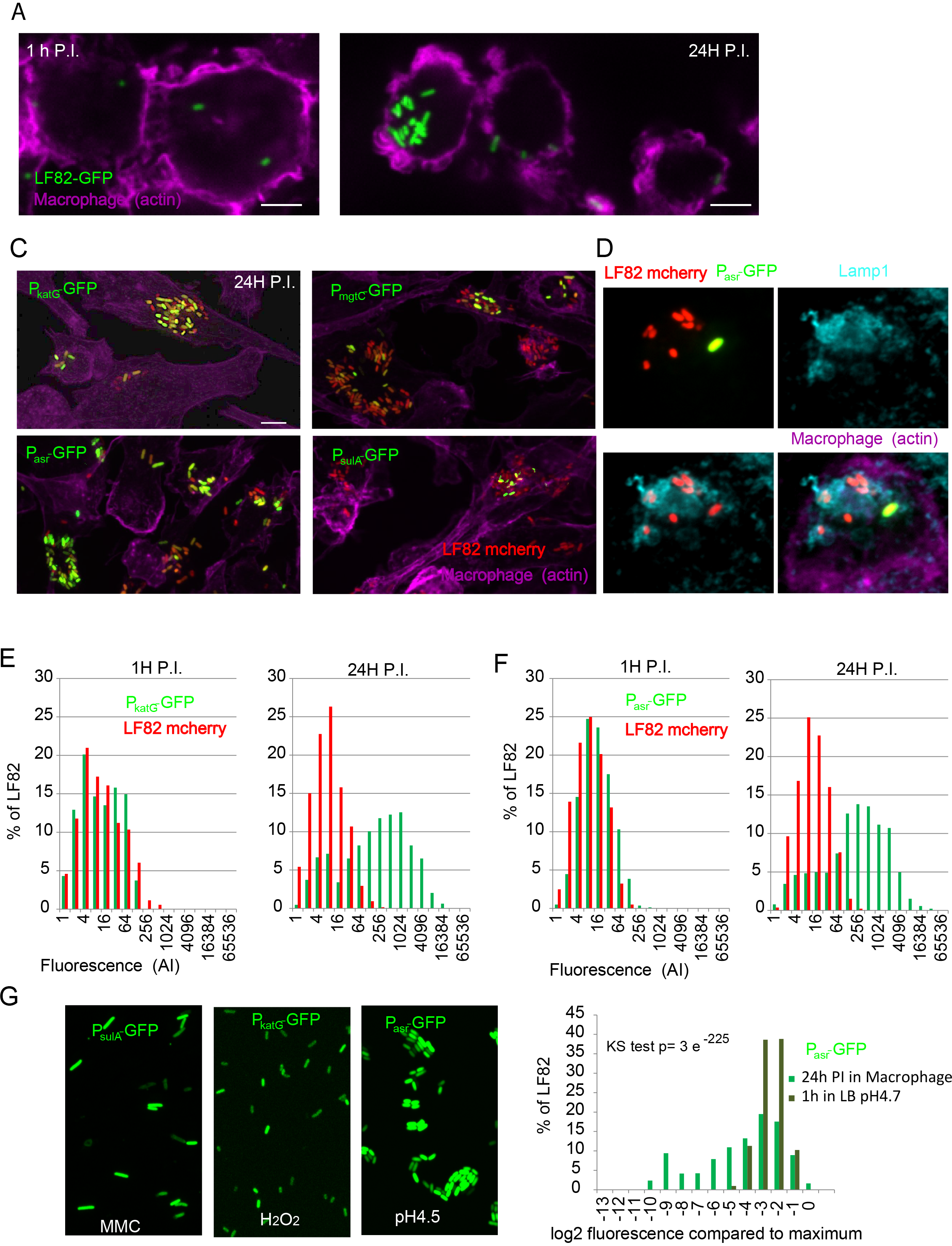
Intracellular LF82 show heterogeneous stress responses. A) Imaging of THP1 macrophages infection by LF82-GFP at a MOI of 30. Representative images at 1 h and 24 h. Scale bar is 5 μm B) Imaging of LF82-mCherry stress responses at the single cell level with biosensors. Imaging was performed at 24 h P.I.. LF82-mCherry was transformed with plasmids containing either the *katG* promoter fused to GFP (P*katG-* GFP), the *mgtC* promoter (P*mgt*C-GFP), the *asr* promoter (P*asr-*GFP) or the *sulA* promoter (P*sulA-*GFP). C) Imaging of LF82-mCherry P*asr*GFP and Lamp1 phagolysosome marker E) Measure of the fluorescence intensity of individual LF82-mCherry containing the *katG* promoter fused to GFP at 1 h and 24 h P.I. F) Measure of the fluorescence intensity of individual LF82-mCherry containing the *asr* promoter fused to GFP at 1 h and 24 h P.I. G) LF82 P*sulA*-GFP, LF82 P*katG*GFP, LF82 P*asr-*GFP in LB respectively supplemented with MMC (5μM), with H2O2 (5μM) or switched to pH4.7 1h before imaging. Distribution of the fluorescence of LF82 P*asr-*GFP after 24h post infection in macrophage (from panel B) and after 1 hours of growth in LB buffered at pH4.7. Fluorescence values were expressed as their log2ratio with the average value of the maximum decile (maximum expression). Distributions were compared with a Two-sample Kolmogorov-Smirnov (KS) test.

### Macrophages induce the formation of non-growing LF82

We analyzed heterogeneity in LF82 cell cycle within macrophages by using two complementary fluorescence assays. First, Fluorescent Dilution (FD) highlights bacteria that have not divided since the time of infection (Figure 3A and Supplementary Figure S4A-C).

**Figure 3.**
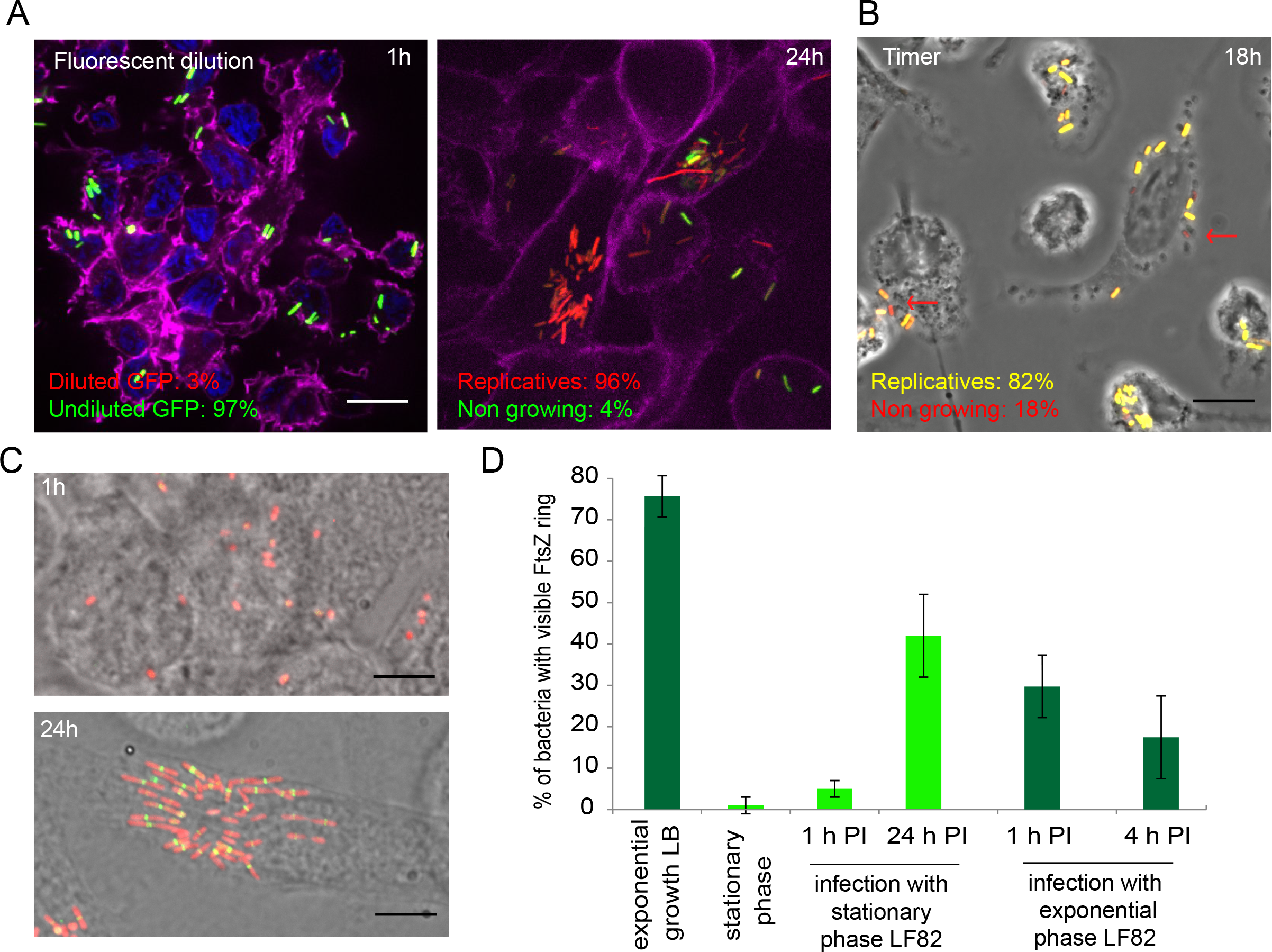
Non-grower LF82 are produced during intracellular growth. A) Representative image of LF82 containing the fluorescent dilution plasmid (pFC6Gi) at 1 h and 24 h post-infection. The frequency of replicative and non-growing LF82 (undiluted GFP) is indicated (N =300). B) Representative image of LF82 containing the TIMER plasmid (pBR-TIMER) at 18 h post-infection. The red arrows points toward the reddest LF82. The frequency of replicative and non-growing LF82 is indicated (N =300). C) Representative images of LF82-mCherry FtsZ-GFP at 1 h and 24 h post-infection. Infections presented in panels A to C were performed with a stationary phase culture of LF82 (O.D. 2). Scale bars are 5 μm. D) Measure of the frequency of LF82 presenting a FtsZ ring in populations growing in LB or within macrophages (N =300).

FD shows that at 24 h P.I. LF82 underwent up to 6 divisions, which is still below the maximum dilution detectable by the assay in our conditions (8 generations in LB, Supplementary Figure S4B). In good agreement with the CFU, measurement of FD showed that only 20% of the population has performed more than 1 division at 6 h P.I. From these observations, we can estimate that the highest generation rate of LF82 within macrophages is ≈ 0.5 doubling /h between 6 and 20 h P.I.. Interestingly, FD also revealed that approximately 4% of the population did not divide or divided fewer than 2 times intracellularly in 24 h (Figure 3A and Supplementary Figure S4B). By contrast among the small amount of K12-C600 bacteria that survived for 24 h in the macrophage, 60% of K12-C600 bacteria underwent fewer than 2 divisions and less than 10% underwent 5 divisions (Supplementary Figure S4D). Second, we used TIMER to refine these observations; it provides an instantaneous evaluation of the generation time during the infection kinetics (Figure 3B and S4A and S4B). TIMER indicated that at 18 h P.I., 18% of the LF82 population was not actively dividing, supporting the existence of a non-growing or slow-growing subpopulation. Since they require dilution of fluorescent proteins both TIMER and FD are poorly informative about the first hours of the infection. Therefore, we used a GFP fusion with the septal ring protein FtsZ to monitor division in the individual bacterium (Figure 3C). In LB, exponentially growing LF82 frequently presented the FtsZ ring (70% of the population), but stationary phase LF82 rarely presented the FtsZ ring (<2% of the population, Figure 3D). Following infection of macrophages with the stationary phase culture of LF82 *ftsZ-gfp*, we observed that 5% (+/−2) and 40% (+/− 12) of the population, respectively, presented a FtsZ ring at 1 h and 24 h P.I. (Figure 3C and 3D). Following infection with exponentially growing LF82 we observed a sudden reduction in the number of LF82 presenting a FtsZ ring at 1 h and 4h P.I. (Figure 3D). The three reporters (FD, TIMER and FtsZ) provided complementary indications: i) within macrophages, LF82 strongly slow down their cell cycle for several hours; ii) starting at 6 h P.I. LF82 multiply; in this phase the generation time may vary among bacteria but can be as short as 2h; iii) a part of the population, completely halted their cell cycle and become non-grower. The difference between the number of non-growing LF82 revealed by TIMER and FD shows that non-growing LF82 are formed late in the infection kinetics and not only upon phagocytosis.

### Macrophages induce the formation of antibiotic-tolerant LF82

Non-growing Salmonella phenotypes have been observed inside macrophages and during mouse infection (Helaine *et al.*, 2014; Claudi *et al.*, 2014). Being tolerant to subsequent antibiotic challenge they were recognized as persisters. We inquired whether also for AIEC LF82 the non-growing component of the population had enhanced antibiotic tolerance. In exponentially growing liquid cultures, approximately 0.01% LF82 tolerated a 3 h ciprofloxacin challenge, and can therefore be considered persisters (Figure 4A). Following a brief passage through the macrophage, the frequency of LF82 bacteria tolerant to ciprofloxacin increased to 0.5 % (nearly 50 fold compared to exponentially growing LF82). Interestingly the number of tolerant LF82 reached 5% (500 fold compared exponentially growing LF82) after 24 h in the macrophage (Figure 4A). This near to 10-fold increase in the number of tolerant bacteria after a 24 h intracellular period compared to a 1 h intracellular period indicates that, like non-growers, tolerant phenotypes are not exclusively formed upon infection, but also as bacteria multiply inside the macrophage. In this respect, the behavior of LF82 differs significantly from *S. typhimurium*, which forms a large number of persisters upon macrophage entry, but this population remains stable during the infection (Helaine *et al.*, 2014). To test if non-growing LF82 revealed by the FD assay indeed corresponded to the antibiotic-tolerant population, we used the macrophage-permeable antibiotic ofloxacin. We added ofloxacin (10x MIC) for 4 h after 20 h of intracellular growth, and we observed significant increases in the proportion of green LF82 (non-growing) inside macrophages (Figure 4B). These observations suggest that the sub-population of non-growing bacteria largely overlaps with that of persisters, where protection from antibiotics may also confer enhanced tolerance to intracellular stresses.

**Figure 4.**
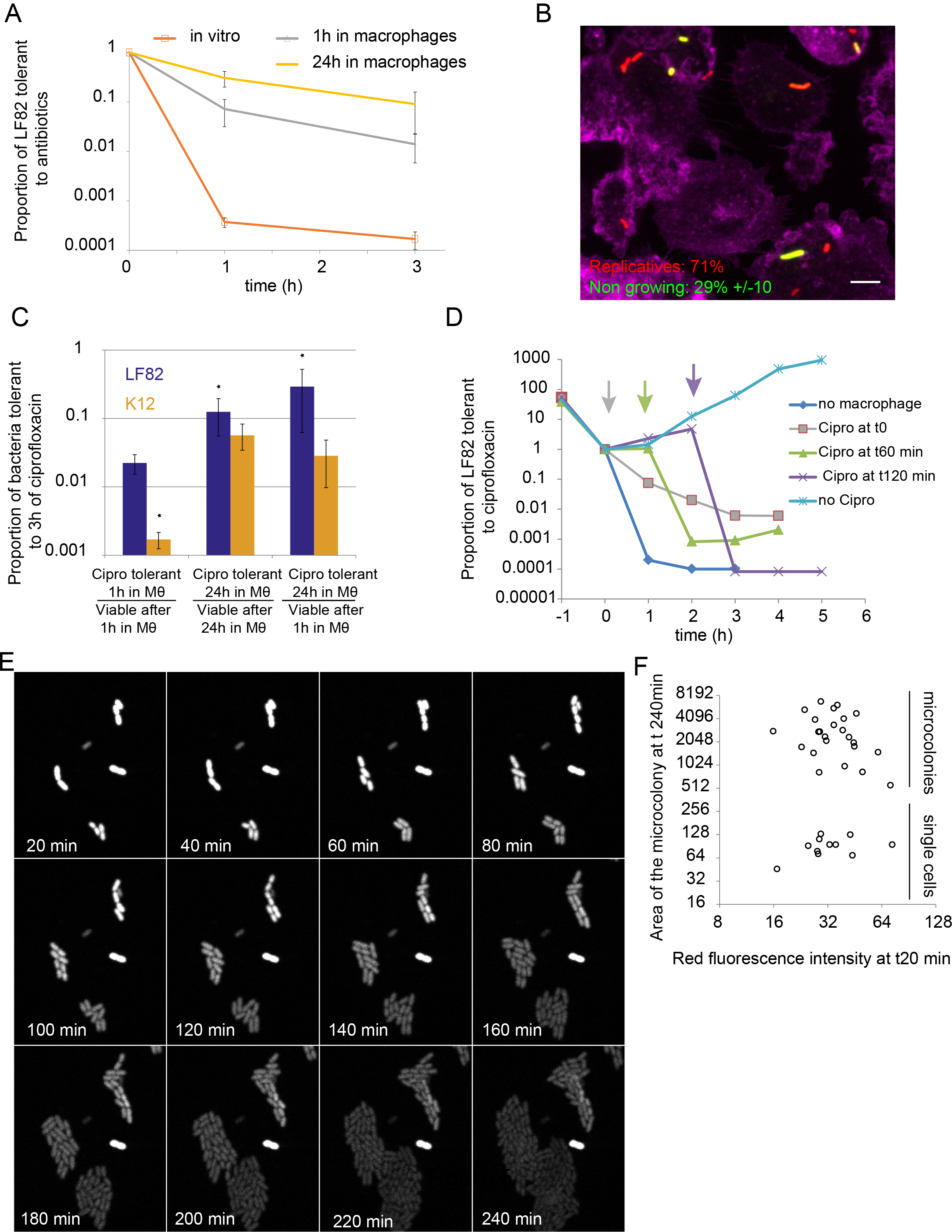
Persisters LF82 are produced during intracellular growth. A) Measure of the proportion of LF82 that were tolerant to ciprofloxacin (10x MIC) at 1 h or 3 h. LF82 were cultivated up to OD 0.3 in LB medium (*in vitro*) or harvested after 1 h or 24 h post infection within macrophages. The challenges exerted on bacteria passaged through macrophages started immediately after macrophage lysis (see experimental procedures). B) Proportion of non-growing LF82 (labeled using the Fluorescent Dilution assay) observed within macrophages (24 h P.I.) following a 6 h ofloxacin treatment. C) Ratio of ciprofloxacin-tolerant versus viable LF82 and K12-C600 bacteria after macrophage infection. Values are averages of 5 experiments. Data were analyzed using a Student’s *t* test to determine differences with the proportion of ciprofloxacin-tolerant LF82 at 1 h post-infection, \**P* < 0.05. D) Measure of the proportion of LF82 that were tolerant to ciprofloxacin at 1 h or 3 h with increasing times after macrophage lysis. E) Imaging of the regrowth properties of individual LF82-TIMER bacteria after macrophages lysis. Infections were performed for 20 h and then macrophages were lysed, LF82 spread on to LB agarose pads and immediately imaged at 37°C. TIMER red fluorescence was progressively lost as microcolonies formed. The lysis procedure requires 20 minutes before the first field can be observed (t20 min). F) Measure of the ability of LF82 to form microcolonies as a function of red TIMER fluorescence at t20 min of the experiment are presented in E; areas of microcolonies are expressed in pixels^2^;

### Tolerance to antibiotics is enhanced for LF82 compared with K12-C600 *E. coli*

We compared the number of LF82 and a non-pathogenic K12-C600 laboratory strain with tolerance to ciprofloxacin following brief (1 h) or long (24 h) passages in macrophages. After a brief passage in macrophages, the proportion of LF82 that were tolerant to ciprofloxacin was significantly higher for LF82 than K12-C600 (Figure 4C). Interestingly, even if the absolute number of ciprofloxacin-tolerant K12-C600 was largely reduced compared with LF82, their proportions among bacteria that survived 24 h inside macrophages were comparable (Figure 4C). These findings demonstrate that the number of antibiotic-tolerant bacteria formed in response to macrophage attack is reinforced for LF82 compared with the laboratory strain.

### Tolerance to antibiotics is a transient state

We next evaluated whether the antibiotic tolerance was a stable or transient phenotype. We used the macrophage lysis procedure to recover LF82 with induced persistence for 1 h in the macrophage; then, we either challenged them immediately with ciprofloxacin or allowed them to recover in LB for 1 h or 2 h before antibiotic challenge. When bacteria were cultured for 1 h in LB, the frequency of tolerant bacteria was decreased in comparison to bacteria that were immediately treated with the antibiotic; however, this number was still higher than that of bacteria that had not infected macrophages. Two hours in LB was sufficient to cause a comparable frequency of ciprofloxacin-tolerant LF82 to that of bacteria that had not encountered macrophages (Figure 4D). These observations show that when the environment is no longer stressful, antibiotic-tolerant, non-growing LF82 rapidly switch back to a replicative mode.

### Characterization of non-growing LF82

Both FD and TIMER revealed slightly more non growing LF82 (4% and 18% respectively, figure 3A) than antibiotic-tolerant LF82 after macrophage lysis (0.5% at 1 h P.I. or 5% at 24 h P.I., Figure 4A). This finding raised the possibility that persisters only form a portion of the non-growing population. To quantify this proportion, we infected macrophages with TIMER-tagged LF82, lysed the macrophages and allowed bacterial growth on a LB-agarose pad under the microscope at 37°C. Seventy percent of the LF82 bacteria recovered quickly from the challenge and formed microcolonies, but approximately 30% of them never divided (Figure 4E). These non-cultivable LF82 presented either non-growing or growing TIMER fluorescence (Figure 4F). The presence of non-cultivable LF82 among the bacteria with non-growing TIMER fluorescence explains the difference between fluorescence and antibiotic assays.

### SOS and stringent responses influence antibiotic tolerance

Among mutants that affected LF82 survival (Figure 1C), only the *recA*, *relA spot* and *dksA* deletions negatively impacted the number of LF82 that were tolerant to a 3h ciprofloxacin treatment (Figure 5A). The impact of the *recA* deletion might be misinterpreted because ciprofloxacin alters DNA and limits resuscitation of *recA* persisters. Therefore, we repeated the tolerance assay with cefotaxime for the following SOS mutants: *recA* (impaired for DNA lesion repair and SOS induction), *lexAind-* (unable to induce SOS) and *sulA* (unable to block cell division). I*n vitro*, SOS mutants did not present defect for cefotaxime tolerance (Supplementary Figure S3B). However, these mutants exhibited a decreased tolerance to antibiotics when persisters were induced by a pretreatment with subinhibitory concentrations of ciprofloxacin (Supplementary Figure S3C). This finding is in good agreement with previous reports (Dörr *et al.*, 2009), and it confirms that SOS induction favors the production of persisters. We analyzed cefotaxime tolerance of these SOS mutants after a 1 h or 20 h period within macrophages (Figure 5B and 5C). We observed a significant reduction of the proportion of *recA* and *sulA* mutants that were tolerant to cefotaxime treatment after a 20 h passage in the macrophage (Figure 5C) but no effect on bacteria that remained only 1 h in macrophages (Figure 5B). The *lexAind-* mutation did not change the number of tolerant bacteria in these conditions. We also analyzed the *dksA* mutant in these assays; surprisingly, it behaved differently than the *recA* and *sulA* mutants: we observed a significant reduction of the proportion of cefotaxime tolerant bacteria following a 1 h passage within macrophages (Figure 5B) but not when the bacteria remained for 20 h in macrophages (Figure 5C). To test an eventual epistatic relation between SOS and stringent response we combined *recA* and *dksA* deletions. They had an additive impact on the ability of LF82 to become tolerant to cefotaxime after a brief infection (Figure 5B) and a 20-hour infection (Figure 5C). Our observations suggest that production of persister/tolerant LF82 bacteria is under the control of the stringent response in the first hours of infection and controlled by genotoxic stress, the SOS response and DNA lesion processing later in the infection. When one of these two responses is deficient the production of persister /tolerant LF82 requires the other.

**Figure 5.**
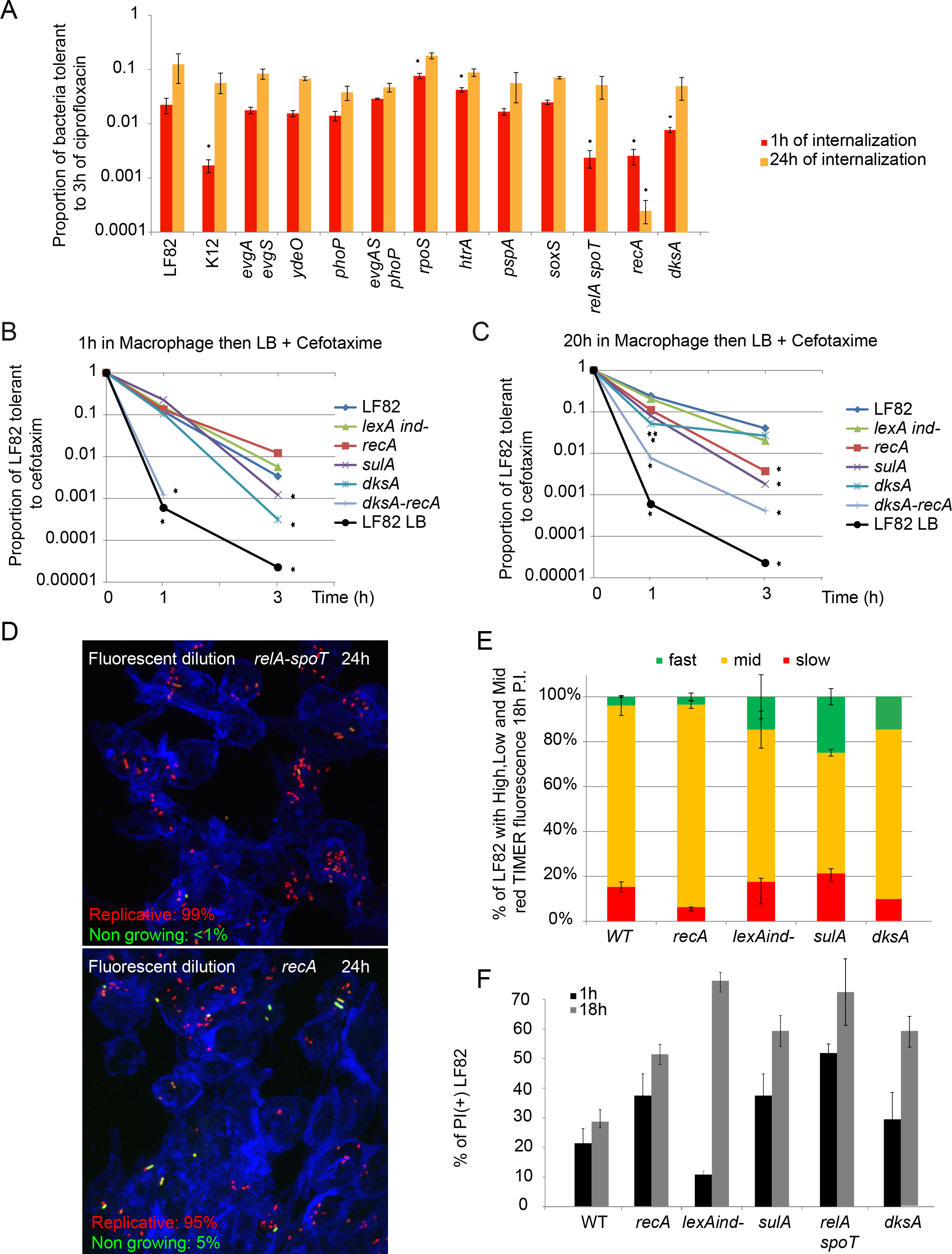
SOS and stringent responses control LF82 cell cycle in the macrophages. A) Proportions of LF82, K12-C600 and LF82 deletion mutants that were tolerant to a 3h ciprofloxacin challenge following a 1 h or 24 h intracellular period within THP1 macrophages. Values represent the average of 3 to 7 experiments. Data were analyzed by using a Student’s *t* test to determine differences compared with wild-type LF82. \**P* < 0.05. B) Measure of the proportions of LF82 *lexA ind*, *recA*, *sulA*, *dksA*, *dksA lexAind-*, *dksA recA* mutants that were tolerant to 3 h of cefotaxim challenge after 1 h of macrophage infection. The 3 htime point was below detection limit for the *dksA recA* mutant because of the poor viability of the mutant in macrophages. Cefotaxime only kills growing bacteria; therefore we resuspended LF82 in LB after macrophage lysis. Under these conditions, we did not observe the plateau observed for persisters to ciprofloxacin, suggesting that tolerant rather than persister bacteria were measured. C) Same as in B but with 20 h of macrophage infection. Values represent the average of 3 to 7 experiments. Data were analyzed using a Student’s *t* test to determine differences compared with wild-type LF82. \**P* < 0.05. D) Imaging of the FD for the *relA spoT* and *recA* mutants. Imaging at 24 h P.I. at MOI 100. Data represent the % of replicative and non-growing LF82 at 18 h; a total of 300 bacteria were counted. E) Percentage of the LF82 population presenting a high, mid or low level of red TIMER fluorescence at 18 h P.I.. The *recA*, *lexAind-*, *sulA* and *dksA* mutants were tested. F) Live and dead assay performed 1 h and 18 h P.I., in the LF82, LF82*recA*, LF82*lexAind-*, LF82*sulA*, LF82*relAspoT* and LF82*dksA* strains.

### The role of SOS response and stringent response for the control of LF82 cell cycle in the macrophages

Knowing that SOS and stringent responses influence the production of antibiotic tolerant LF82 after a passage within macrophage we examined whether they also contributed to LF82 cell cycle control, i.e. production of non-growing, replicative or dead LF82. We used the FD assay to measure the number of non-growing LF82 in *the relA-spoT* and *recA* mutants. FD revealed that the non-grower number was dramatically reduced in the *relA-spoT* mutant (<1%) (Figure 5D). This suggests that in the absence of stringent response LF82 cannot immediately curb its cell cycle upon phagocytosis. By contrast the number of non-growers was unchanged for the *recA* mutant (Figure 5E). This is in agreement with the absence of effect of the *recA* deletion on the production of cefotaxime tolerant LF82 early in the infections. TIMER revealed that at 20 h P.I. the proportion of slow, mid and fast growing LF82 was affected by the alteration of *recA, lexAind-, sulA* and *dksA.* The *recA* and *dksA* deletions reduced the number of non-growing LF82; by contrast, the *lexAind-* and *sulA* mutations only increased the number of fast growing LF82 in the population (Figure 5F). Finally, the live and dead assay showed a strong increase in lethality of the *recA, sulA*, *lexAind-, relA spoT* and *dksA* mutants at both time points (Figure 5G) suggesting that in the macrophage environment a failing cell cycle control will almost certainly lead to LF82 death.

Altogether our results showed that the stringent response is the main controller of the early intracellular survival of LF82; it limits LF82 growth and induces the formation of non-growers and among them persisters. Later on, when replication is resumed, SOS response grows in importance. DNA lesions that have been accumulated in the lag phase must be repaired to allow replication and formation of new non-growers and new persisters.

### Kinetics of macrophage infection

AIEC LF82 tolerates macrophage induced stresses, thus it survives and multiplies in the phagolysosome. The population expansion is accompanied by a rise in the number of bacteria that do not grow and tolerate an antibiotic challenge (called henceforth persisters). The change in time, during macrophage infection, of LF82 population size and fraction of persisters are relevant to future fundamental studies, but also to devising therapeutic strategies involving AIEC or other intracellular pathogens. In order to explore the mechanistic bases of the infection kinetics, we have used a mathematical model (Figure 6A) to fit the observed changes in CFU and persister counts during 24 h for LF82, K12-C600 and the stringent response mutant LF82*dksA* (Figure 6B). The model is based on the following biologically-informed hypotheses (illustrated in Figure 6A and detailed in the Supplementary text 1). i) Reproduction: the population of replicating bacteria B has a constant net growth rate (birth minus death rate) δ_1_, which is either 0 or negative during a lag phase of duration λ, and β>0 otherwise. ii) A stress-induced death rate, δ_2_(S) that increases with stress (S). We assume that stress S, possibly due to lesions (DNA lesions, membrane alterations, oxidative damages among others) that accumulate over time or due to a macrophage response to the infection, builds up in time proportionally to the total number of bacteria in the population. iii) Switch to persistence: bacteria have a constant probability k_p_ of generating non-growing, stress-tolerant phenotypes P. The dynamics of a population of bacteria can be described by a set of three ordinary differential equations for the number B of non-persister bacteria, the number P of persisters, and the stress variable S (Methods). The number of dead bacteria D can be derived from these under the assumption that dead LF82 bacteria decay exponentially with rate 0.56 (computed from the assay in Supplementary Figure 2C). The model has a total of 12 parameters for the three strains, which are fitted to data as explained in Methods and SI. Although LF82 displays a considerable overshoot in population size (as also observed in (Glasser *et al*, 2001)), the dynamics can be reproduced by choosing the same β for every strain. We estimated such net growth rate to 0.15 ± 0.003 h^−1^, corresponding to 0.21 divisions per hour (Figure 6B), consistent with independent cell-level measures by FD (Supplementary Figure 4). The most notable quantitative difference between strains is that K12-C600 displayed a lag phase of more than 13 h, twice as long as LF82 and LF82*dksA*. A consequence of this difference is that when K12-C600 bacteria start actively duplicating, stress has already built up. Together with K12-C600’s enhanced sensitivity to stress, this curbs the population expansion, resulting in a lower overall growth within macrophages. In the LF82*dksA* mutant, growth is instead impaired by increased initial mortality (whose rate δ_1_ has been estimated by PI measures, Supplementary Figure S2C), presumably related to stringent response failure. With respect to the other strains, moreover, LF82 is advantaged at later times – when the SOS response becomes important – thanks to reduced stress-induced death rate. Rate of persistence production for LF82 and LF82 *dksA* (0.08 and 0.002 h^−1^, respectively) is estimated to be higher than for K12-C600 (0.001 h^−1^), supporting the notion that AIEC strains within macrophages turn to persisters at an enhanced rate, but less so if their stringent response is impaired. The model allows testing changes in infection dynamics for ‘virtual mutants’ LF82*, obtained by varying k_p_, λ and d_max_ – the parameters that quantitatively differ between LF82 and *E. coli* K12-C600 (Figure 6C). The total population overshoot is enhanced when lag phase is shorter and the effect of stress less acute, but damped when persisters production is more frequent. Interestingly, the phenotypes of the single stress response mutants (acid, oxidative, lack of Mg^2+^), i.e. reduced CFU at 24 h without perturbation of the persister proportion in the population (Figure 1C and 3A) were nicely reproduced by a change in the single d_max_ parameter. This suggests that the model can be used to plan future works on the effect of mutants or drugs on macrophage colonization by AIEC.

**Figure 6.**
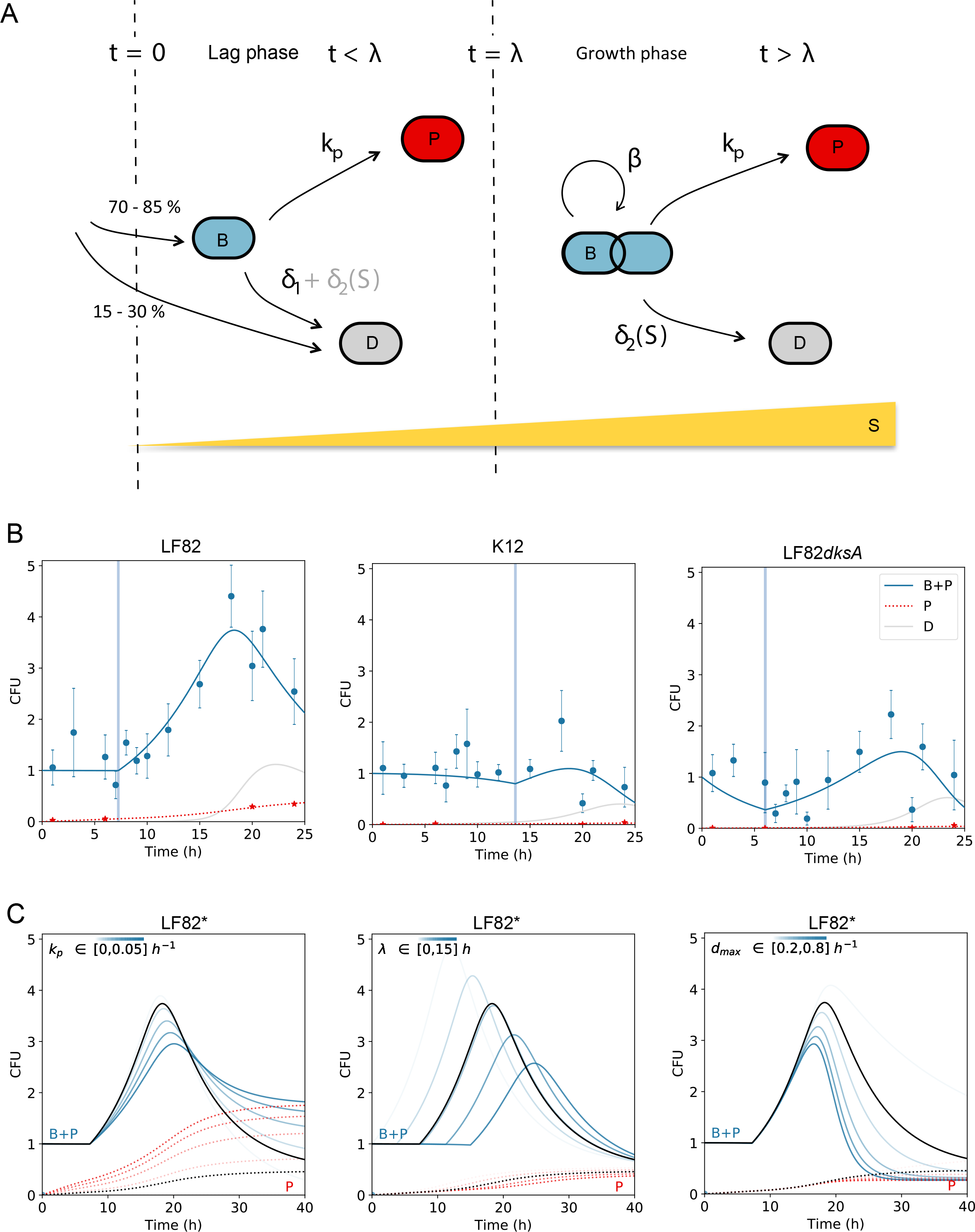
Kinetics of macrophage infection. A) Model of infection of THP1 macrophages by LF82, describing the processes of net growth, switch to persistence and stress-induced death as explained in the text. Part of the cells that enter the macrophage die at the onset of the infection (t=0). During lag phase (0<t<λ), death either exactly compensates birth or, in the mutant lacking the stringent response, results in a negative net growth rate −δ_1_. Later in the infection, the net growth rate β is positive. Death rate due to stress accumulation (yellow bar) is negligible in the early stages of infection and becomes particularly important at late time points. Bacteria switch to a persistent state at a rate k_p_ independent of the growth stage. B) Experimental measures of the infection kinetics (CFUs from 5 replicate experiments, circles; persister fractions, stars) collected over 24h for LF82, *E. coli* K12-C600 and LF82 *dksA* and the best fitting parameters (Supplementary text Table 1) of model eq. (1) (Methods). Continuous lines represent total number of bacteria (B+P, continuous line) and persiters (P, dotted line). Vertical lines indicate the duration λ of lag phase. C) Projected changes in infection dynamics for ‘virtual mutants’ LF82*, obtained by varying k_p_, λ and d_max_ – the parameters that quantitatively differ between LF82 and *E. coli* K12-C600 – around the LF82 best fit solution (black line); colored lines correspond to parameter values within the indicated interval.

## Discussion

We analyzed the growth and survival strategies used by the prototype AIEC strain LF82 to colonize the human monocytes-derived THP1 macrophages. Our analysis revealed that intracellular LF82 are constantly under stress while colonizing macrophages. The consequences of these stresses are important: increase in the death rate of bacteria, slow multiplication of replicating bacteria and formation of a large number of non-growing bacteria. LF82 adapts to this environment thanks to successive phenotypic switches that require the two main stress responses: the SOS response and the stringent response (Figure 7).

**Figure 7:**
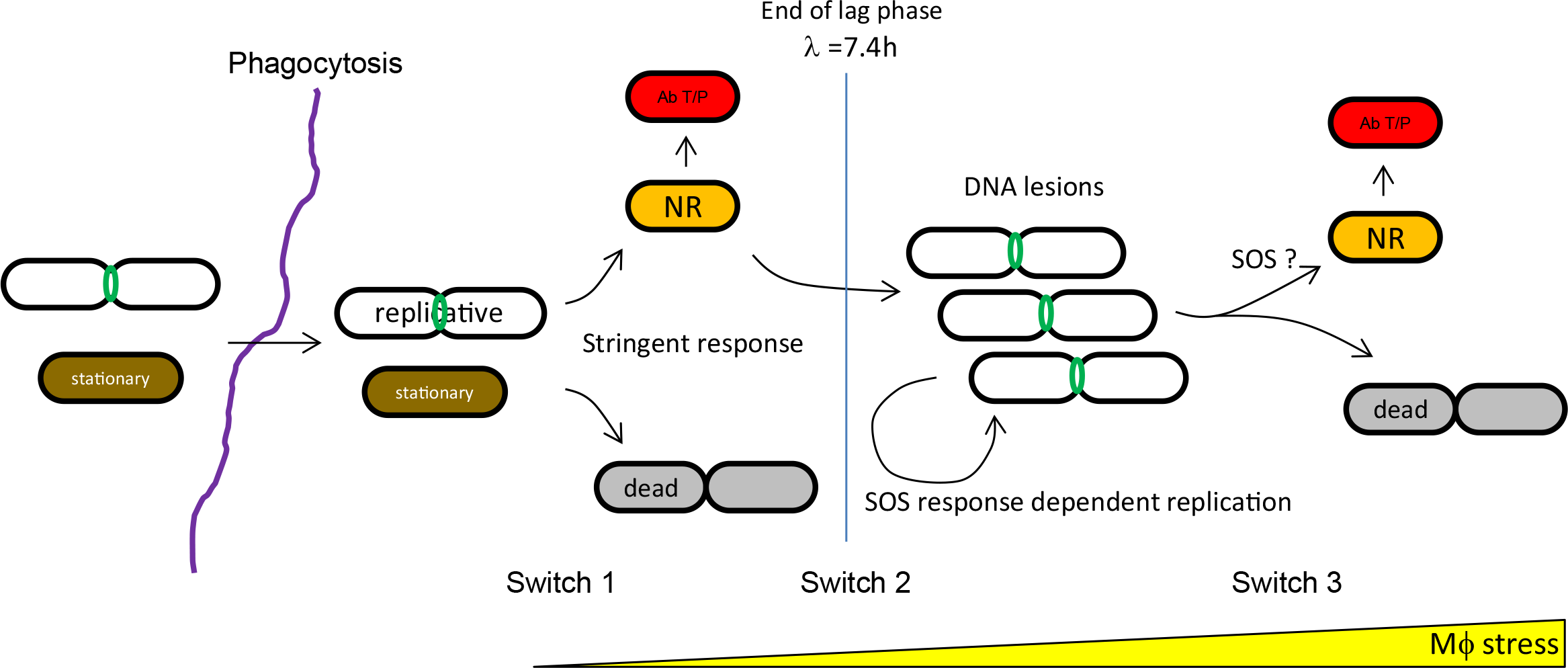
Dynamic of AIEC LF82 bacterial population during macrophage infection. Upon phagocytosis both replicative (green FtsZ ring) or stationary phase (brown) LF82 detect a signal, perhaps nutrient depletion, that led to stringent response activation. This activates a first phenotypic switch toward a non-replicating state (orange) that protects LF82 from dying because of initial stress burst. Among these non-replicating LF82 persisters are formed. After this lag phase, a second switch is required to initiate few rounds of replication. The timing and perhaps the frequency of switching from lag phase to replicative phase differentiate LF82 from our control commensal strain. We have not yet identified LF82 specific determinants that allow this switch. Subsequent replication rounds are dependent on the DNA repair machinery. A third switch, linked to the increasing stress or lesions, is turned on in a portion of the replicative population to form new non growers and persisters. The SOS response might also be playing a role at this stage.

### Macrophages place LF82 under lethal stress

Using fluorescent reporters, we measured that half of the LF82 population present at 24 h P.I. had given rise to 6 or more generations. Under controlled laboratory conditions, this should produce a 30 to 60-fold increase in the population size at 20 h compared with the 1 h time point PI. However, we only observed a 3 to 6-fold increase in viable bacteria at 20-24 h compared with 1 h. We demonstrated that this modest colonization of macrophages by LF82 is explained by a big death rate and switch from replicative to non-growing cell cycle. At the single-macrophage and single-bacterium level, FD and TIMER fluorescent reporters revealed non-growing LF82. We observed macrophages containing few (less than 4) LF82 with red fluorescent dilution staining. These bacteria had therefore divided several times (>4) before observation and thus should be accompanied by their siblings (>16). This observation is in good agreement with our Live and Dead assay indicating that LF82 progeny has a significant chance to be killed and destroyed by the macrophage. By contrast, some macrophages contained growing LF82 and ultimately acquired more than 50 bacteria in one or several compartments. The live and dead assay confirmed that LF82 was frequently killed by macrophages. Because alterations of bacterial stress responses significantly reduced the bacterial yield, we propose that LF82 death is the consequence of oxidative, acid, genotoxic and protein stresses imposed by the macrophage. We compounded these experimental observations in a mathematical model describing the dynamics of bacterial infection within macrophages. A first phase of stalled growth, a likely combined effect of a prolonged lag phase and of compensation between death and division, is followed by an exponential increase in bacterial concentration. This second phase might correlate with a transient increased permissiveness of phagolysosomes or more likely the adaptation of LF82 to growth in this stressful environment. This expansion is successively curbed by the building-up of stressors. Many persisters are formed in the first phase. However, in the second phase a phenotypic switch to non-growing LF82 will eventually result in a sizeable increase of the persister population. We thus understand the survival of LF82 as a consequence of its ability to adapt to harsh phagolysosome environment both at the entry of the macrophage, by induction of stress responses and particularly the stringent response, and during exponential expansion by SOS response. LF82 advantage over K12-C600 would reside in its ability to exit from the lag phase to perform a few rounds of replication/division before stress becomes too strong (Figure 6). Strategically, early onset of growth is compensated by production of persistent bacteria, which endows the pathogenic strain with long-term survival in spite of rapid exploitation of the macrophage environment. We have not yet identified LF82 specific regulons, genes or mutations that allow this transition to take place.

### Replicative LF82

Fluorescent dilution revealed that after the exit of lag phase LF82 replicated moderately within macrophages, with generation time longer than 2 h. *In vitro*, this would be comparable with generation times observed in minimal medium with poor carbon sources such as acetate. Our observations revealed that within macrophages, 40% of the LF82 population presented an FtsZ ring, which is significantly above the number expected from a mixed population of *E. coli* growing with a 2h generation time (28% of cells with FtsZ ring (den Blaauwen *et al*, 2001)) and non-growing cells. Interestingly, in spite of SulA induction, we did not observe filamentation of LF82 within macrophage. These finding demonstrates that some of the cell cycle rules that were established under defined *in vitro* conditions do not apply to intracellular growth conditions, opening avenues for future investigations of bacterial cell cycle regulation in the context of host infection or the microbiota.

### Non-replicative LF82

FD and TIMER revealed that a significant number of intracellular LF82 were not growing. FD revealed that approximately 4% of the phagocytosed LF82 immediately halted their cell cycle. TIMER revealed that at 20 h P.I., approximately 20% of the LF82 population was not growing. We also demonstrated that once the macrophages were lysed, a large portion of the LF82 population (from 0.3 to 10%) was tolerant to several hours of antibiotic challenge, and the proportion of non-growers in the population increased in macrophages in the presence of antibiotics. Altogether, these observations suggest that the phagolysosome environment induces frequent cell cycle arrests among the population and that a part of this arrested population is persisters or tolerant to antibiotics. Such a phenomenon has been previously described during *S. typhimurium* infection of macrophages (Helaine *et al.*, 2014) or mice (Helaine *et al.*, 2014; Claudi *et al.*, 2014) and is reminiscent of VBNR mycobacteria (Manina *et al.*, 2015). Interestingly, we observed an increase in the proportion of macrophage-induced antibiotic-tolerant LF82 at later time points, suggesting adaptive responses to the intracellular microenvironment.

### Stress responses are important for LF82 survival within macrophages

As expected for bacteria residing in a toxic environment, stress responses are important for LF82 survival within macrophages. Acidic, oxidative, genotoxic, and envelope alterations, lack of Mg2+ and lack of nutrient stress responses significantly decreased the fitness of LF82 (50 to 10% of WT). In a few cases, we demonstrated an additive effect of simultaneously altering two pathways. However, LF82 demonstrated surprisingly good tolerance to these alterations compared with the *in vitro* findings for individual stresses. For example, *recA* deletion mutant was extremely sensitive (<1% survival) to prolonged treatment with genotoxic drugs (Supplementary Figure S3A); by comparison, in macrophages, despite clear SOS induction, the viability of the *recA* mutant was only reduced by half compared with wild type bacteria. Set aside the possibility that stress-less niches exist due a possible heterogeneity in the macrophage population, the fitness decline of LF82 stress mutants may be limited by a combination of slow growth, formation of non-growers and/or yet uncharacterized adaptation pathways.

### SOS and stringent responses successively control LF82 fate

We investigated the trigger that could allow some LF82 to halt their cell cycle inside macrophages. It appeared to be unrelated to the ability to sense acidic or oxidative stress. At an early time point, stringent response mutants altered the survival and production of antibiotic-tolerant LF82 suggesting that abrupt nutrient starvation is one of the first signals received by LF82 upon phagocytosis. The early stringent response should result in a slowdown of transcription, translation and DNA replication, and therefore, it might provoke the formation of non-growers and a 7-h lag phase. We assessed whether the stringent response impaired the declines in viability in this first period, and we observed a decrease in the number of bacteria with undiluted GFP using the FD assay, as well as a reduction in antibiotic-tolerant LF82 induction. This suggests that the slowing down induced by the stringent response confers a temporary protection than can be extended to antibiotic tolerance when bacteria become persisters. Accordingly the impact of stringent response alteration was less apparent after the lag phase when replication is re-established in a portion of the population of LF82. SOS induction was moderate at 1 h but important at 6 h and 24 h P.I.; this is in good agreement with the lack of an effect of *recA* deletion on the accumulation of LF82 with undiluted FD GFP and the lack of an influence of SOS mutants on the number of LF82 that were tolerant to cefotaxim at 1 h after infection. DNA lesions could, however, form during this period, but they were mostly observed when DNA replication restarted after 6 h. In this second phase of infection, SOS induction in replicative bacteria might play several roles: i) sustaining DNA repair and therefore DNA replication, cell division and increases in population size; ii) decelerating the division progression, this is the role of SulA, and thus contributing to the formation of new non-growers; iii) intervening for resuscitation of non-growers presenting DNA lesions, both within macrophages and after macrophage lysis.

### Macrophages as a niche for LF82 survival

The purpose of macrophage colonization by LF82 in Crohn’s disease patients is not yet understood. *In vitro*, LF82 colonization did not provoke extensive death of macrophages, which are thus unlikely to serve as a transient replicative niche for ileal infection. Alternatively, we can imagine that dormant LF82 within macrophages can serve as a long-term storage. In this environment, bacteria might be protected from competition with other species of the microbiota and coincidentally from antibiotics. Upon macrophage lysis or inactivation, dormant LF82 would be released and would start to multiply under adequate conditions.

## Methods

### Strains and plasmids

Deletion mutants (Supplementary Table S1) were constructed using the recombineering method as described in (Demarre *et al.*, 2017). Plasmids are described in Supplementary Table S2.

### Infection and microscopy

THP1 monocytes (5×10^5^ cells/ml) differentiated into macrophages for 18 h in phorbol 12-myristate 13-acetate (PMA, 20 ng/ml) were infected and imaged as previously described (Demarre *et al.*, 2017). Infections were performed at MOI 30 (measured by CFU), resulting in the observation of 3 LF82 bacteria per macrophage on average at 1 h P.I.. Imaging was performed on an inverted Zeiss Axio Imager with a spinning disk CSU W1 (Yokogawa).

### Antibiotic challenge and viable bacterial count using the gentamycin protection assay

To determine the number of intracellular bacteria after 20 min of infection, infected macrophages were washed twice with PBS, and fresh cell culture medium containing 20 μg ml^−1^ of gentamicin (Gm) was added for the indicated time (1 h to 30 h). Cell monolayers were washed once with PBS, and 0.5 ml of 1% Triton X-100 in 1x PBS was added to each well for 5 min to lyse eukaryotic cells (Bringer *et al.*, 2006). Samples were mixed, diluted and plated on LB agar plates to determine the number of colony-forming units (CFU) recovered from the lysed monolayers. For the antibiotic tolerance assays, macrophage lysates were transferred to 5-ml tubes and centrifuged for 10 min at 4100 g. The pellet was either suspended in 1x PBS (t0) and ciprofloxacin (1 μg/ml) for 1 h and 3 h, or in LB and cefotaxim (100 μg/ml) for 1 h and 3 h. CFU were measured by serial dilution. Tolerance was estimated for the 1-h and 3-h time points as a function of the CFU at t0.

### Live and dead assay

At the indicated time points, macrophages were lysed with vigorous resuspension in 1x PBS 1% Triton. The cell lysate was pelleted at 300 g for 10 min to eliminate large cell remnants. The supernatant was centrifuged at 4000 g for 10 min. The bacterial pellet was suspended in 1x PBS and processed with the Live and Dead BacLight Viability kit (Thermo Fisher). The bacteria were pelleted for 3 min at 5000 g, resuspended in 50 μl of 1x PBS and spread on a 1% agarose 1x PBS pad for immediate observation.

### Measurement of gene expression by RT-qPCR

Total RNA was extracted with TRIzol reagent from 10^6^ macrophages, as described in the *Molecular Cloning, a laboratory Manual* (Green and Sambrook, CSH Press). First-strand cDNA synthesis was performed with the Maxima First Strand cDNA Synthesis Kit for RT-qPCR (Thermo Fisher), and real-time qPCR was performed with SYBR Green Master Mix (Bio-Rad) on a MyiQ real-time qPCR machine (Bio-Rad).

### Fluorescence quantification

Custom-made FIJI macros were developed for the analyses of fluorescence. For the Biosensors and FD, constitutive expression of mCherry from p-mCherry was used to construct bacterial masks, which were subsequently used to measure GFP intensity. For TIMER analyses, green fluorescence was used to construct the mask. Fluorescence distributions were analyzed with the distribution fitting tool in MATLAB.

### Mathematical model for the infection kinetics

A set of three ODEs recapitulates the main features of the observed growth within macrophages, as explained in the main text and in the SI:

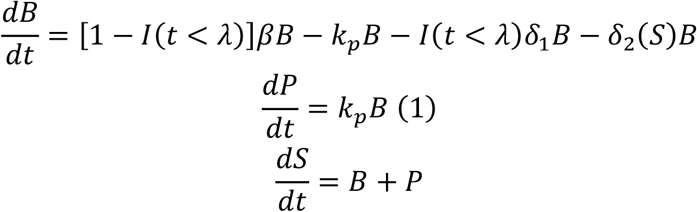

Here, *I*(*t* < λ)is the indicator function, which is unitary during lag phase. At the beginning of the infection, thus, the net growth rate -δ_1_ is either zero (K12-C600 and LF82) or negative (stringent response mutant LF82dskA). At later times, net growth rate β is instead positive. The stress-induced death rate has been chosen to be a sigmoidal function of the stress level:

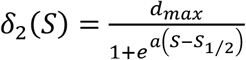

where the half-saturation stress value S_1/2_ and the sensitivity parameter a are assumed to be identical for all strains. Here stress is an effective variable quantifying the effect of crowding on growth within macrophages, and could correspond both to density-dependent reduction of bacterial growth rate (e.g. due to resource depletion), and to the progressive buildup of macrophage-induced killing.

### Fit of the infection kinetics data

Parameters providing the best fit of eqs. (1) to the times series of CFUs and persisters have been obtained by a weighted least-square distance minimization using the python differential evolution algorithm. We used a two-step approach to the fit which allowed us to establish first a subset of 7 parameters (λ and k_p_ for each strain and β) that shape the lag and exponential phases of growth. Subsequently, we fixed β and the λs, and fitted the remaining parameters. Details of the fitting procedure are found in the SI, and the results of the fit in Table 1 of the Supplementary text.

## Supporting information

supplementary informations

## Acknowledgments

We gratefully acknowledge Laurent Aussel, Dirk Bumann, Sophie Helaine, Jakob Moller Jensen and Fabai Wu for providing the biosensors and fluorescent cell cycle reporters. We thank Parul Singh and Xavier de Bolle for careful reading of the manuscript and fruitful discussions. We are very grateful to the members of the CIRB imaging facility. This work has received support from the program «Investissements d’Avenir » launched by the French Government and implemented by ANR with the references ANR--10--LABX--54 MEMOLIFE and ANR--11--IDEX--0001--02 PSL* Research University, from the ANR with the reference ANR-18-CE35-0007 and the support of the association François Aupetit (AFA).

---

**Table S1.**
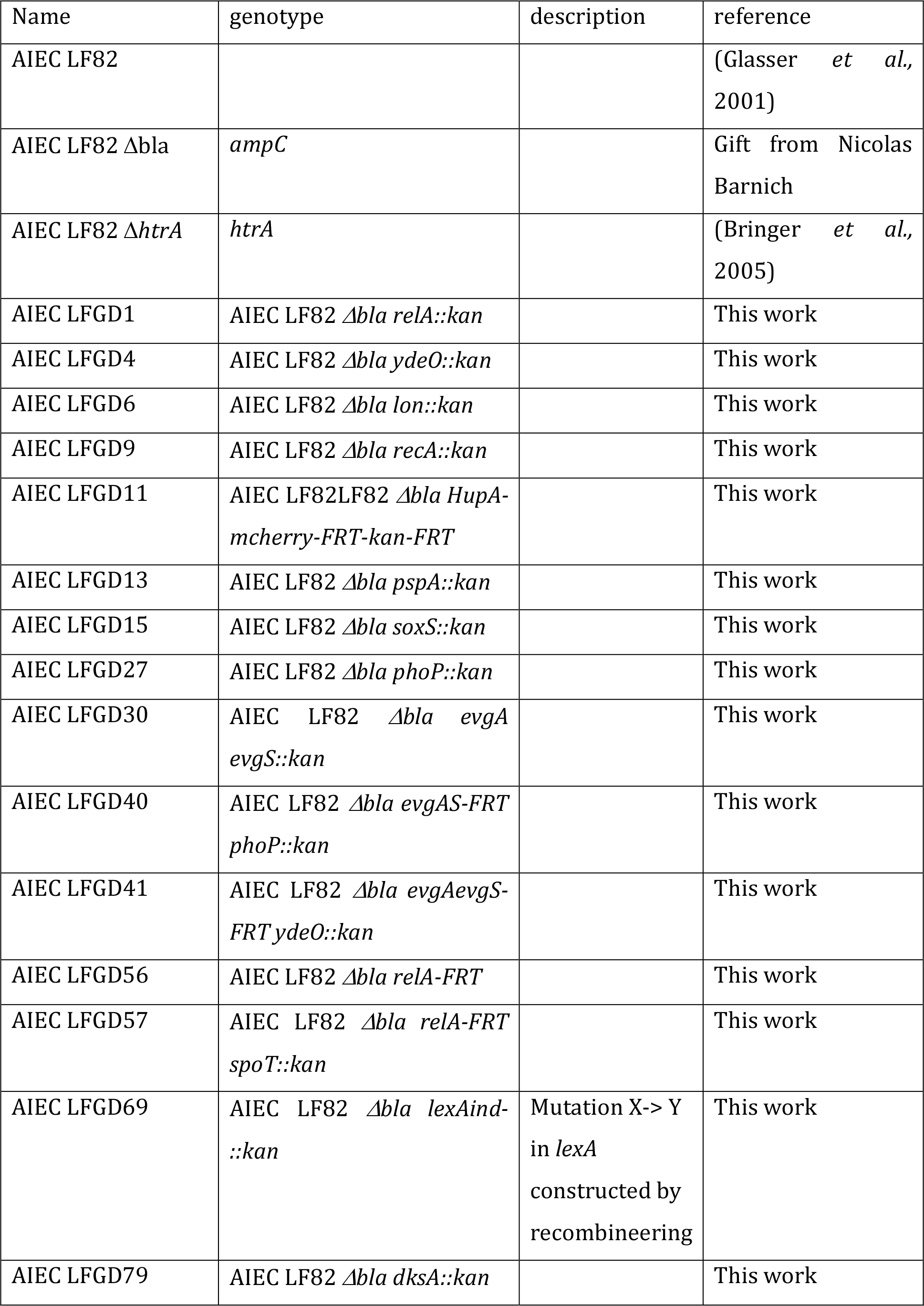
Strains

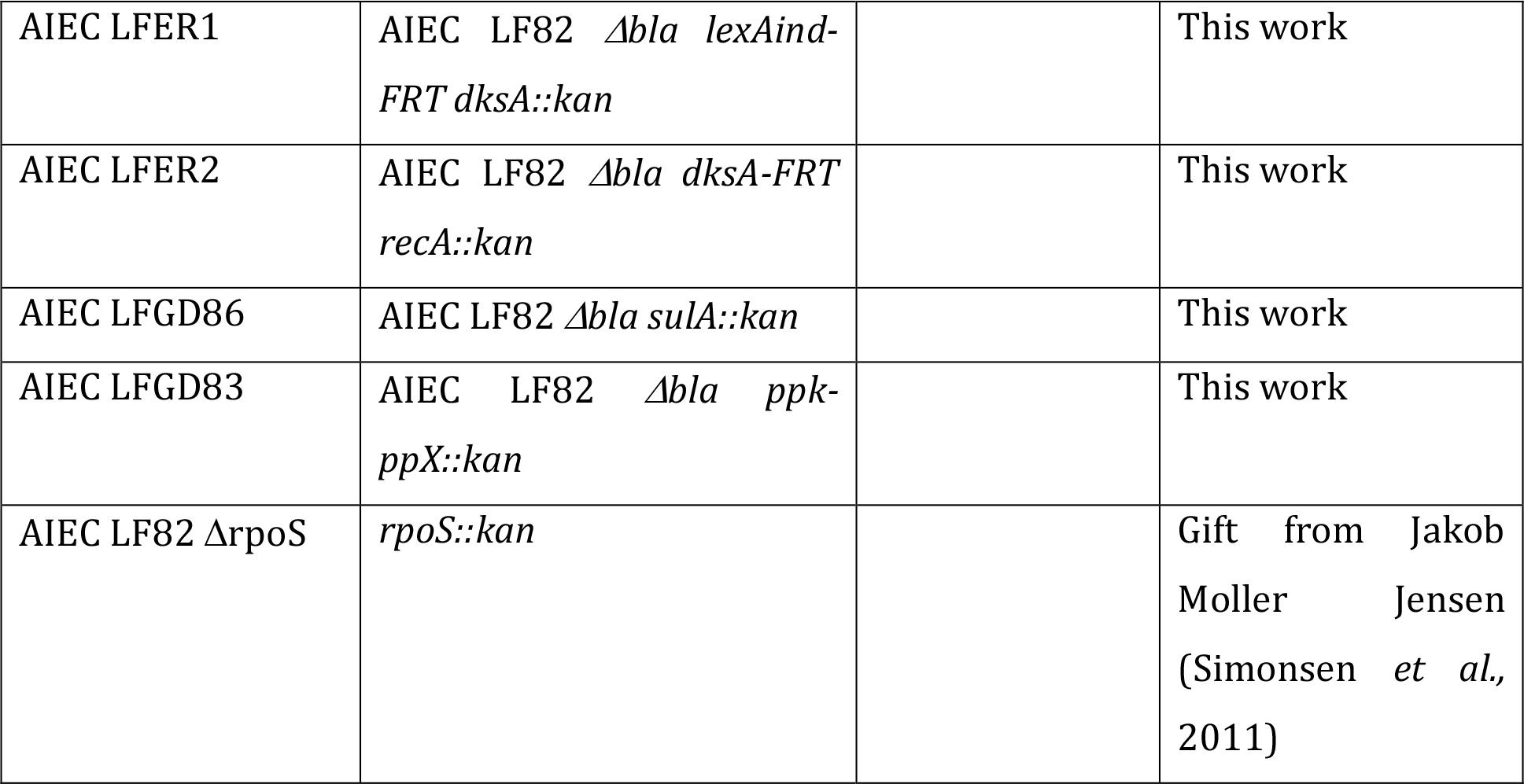

**Table S2.**
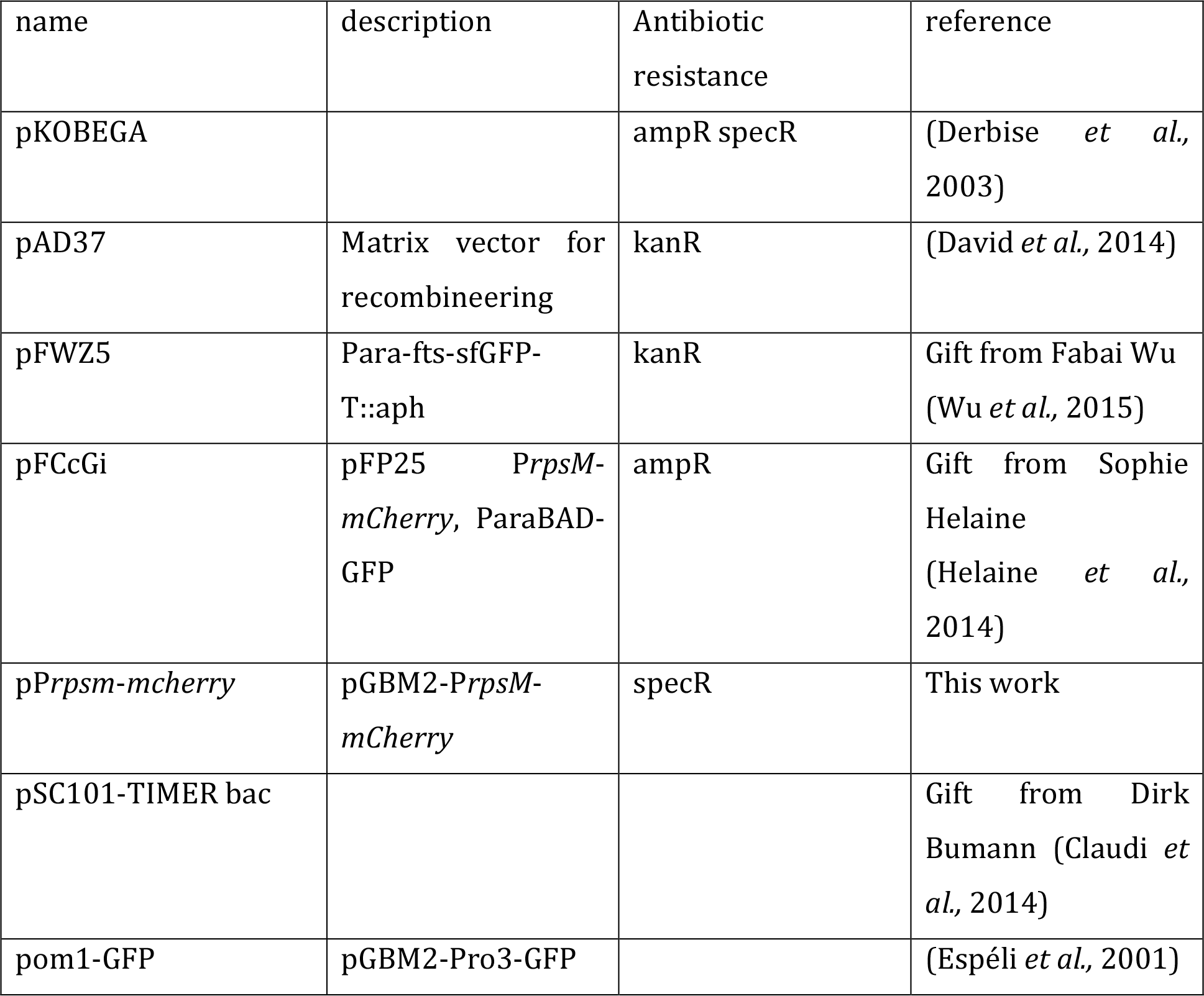
Plasmids

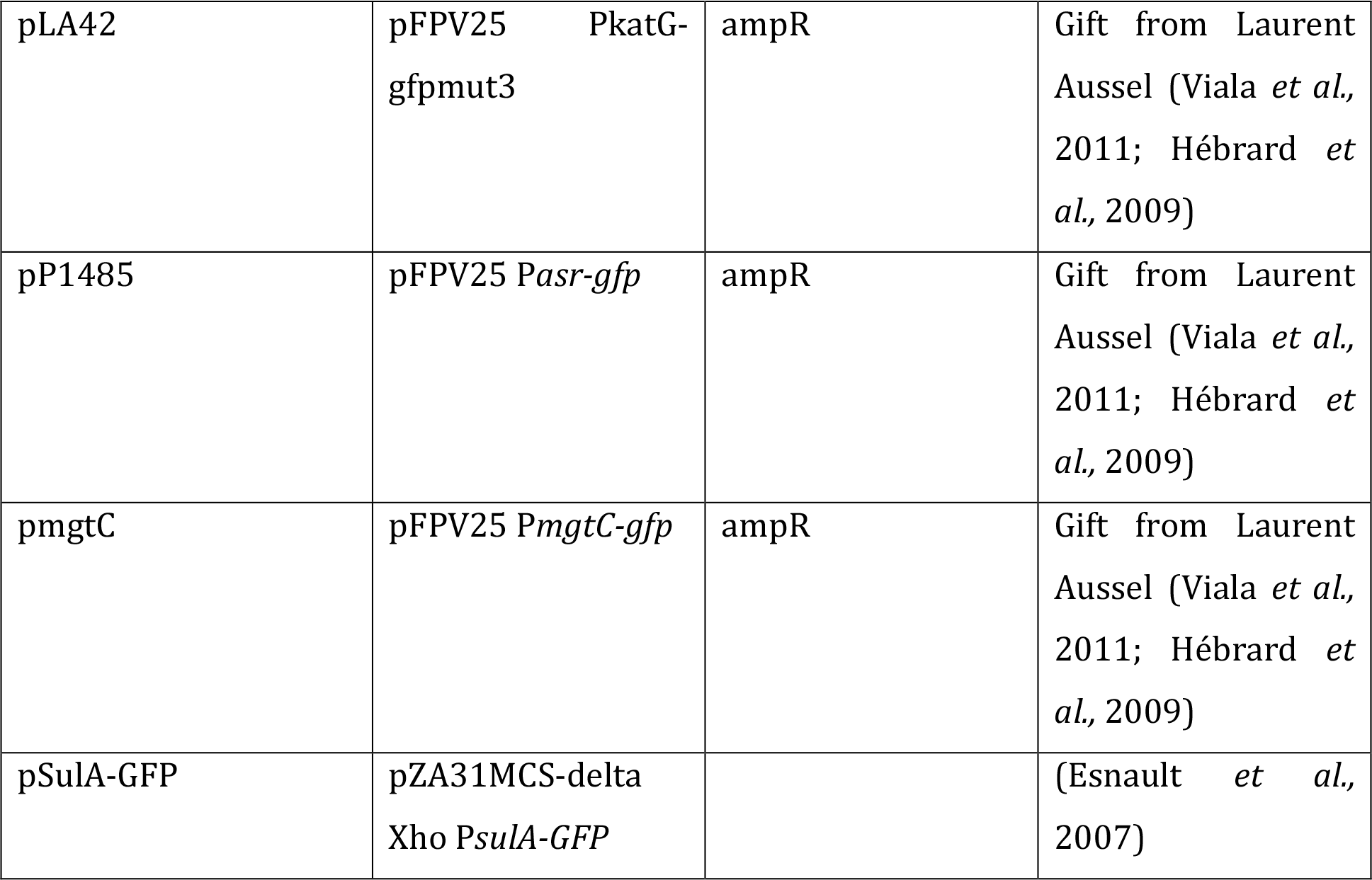

## Legend of the supplementary figures

**Supplementary Figure S1. Measure of the impact of various stresses on LF82 and K12-C600 growth.** A) Growth curves of LF82 and K12-C600 in LB medium at pH 7.4 (blue) and in the presence of serine hydroxamate (15 mg/ml), Serine hydroxamate (7 mg/ml), EDTA (70 mM) or in LB at pH 4.7 and LB at pH 4.7 in the presence of EDTA (70 mM). B) Chemicals or a pH shift were applied at 160 min. B) Same as in A with addition of the antibiotics ciprofloxacin (24 ng/ml) or cefotaxim (800 ng/ml). Data are the mean of 3 technical replicates.

**Supplementary Figure S2. Intracellular Live and dead assays.** A) Live and dead assay performed in situ on infected macrophages or after macrophage lysis. For in situ experiments a very weak propidium iodide (PI) labeling is observed on putative dead LF82. By contrast strong PI labeling is observed after macrophage lysis. B) Measure of the speed of disappearance after phagocytosis by macrophages of heat-killed LF82. LF82 were killed by 15 min incubation at 60°C and subsequently labeled with propidium iodide. Labeled dead LF82 were incubated with macrophage at a MOI of 100. Imaging was performed at 1h, 2h, 3h and 24h post infection. SYTO-9 was used to reveal macrophage and eventual live bacteria. The number of dead LF82 per macrophage was measured at each time points. Data are average of 3 experiments.

**Supplementary Figure S3. Impact of SOS mutations on survival and persistence of LF82 strains.** A) Measure of the resistance to Mitomycin C (MMC) of LF82, LF82 *recA*, LF82 *lexAind-*, MG1655, MG1655 *recA* and MG1655 *lexAind-*. B) Measure of the tolerance to cefotaxim of LF82 and LF82 *recA*, LF82 *lexAind-*, LF82 *sulA* grown in LB medium to an OD of 0.2. C) Induction of tolerance to cefotaxim by pretreatment with a subinhibitory dose of ciprofloxacin (24 ng/ml). The data represent the ratio of the number of bacteria that were tolerant to 3 h of cefotaxim in the presence or absence of ciprofloxacin.

**Supplementary Figure S4. Calibration of the fluorescent dilution assay.** A) Growth curve of LF82 pFC6Gi in LB at 37°C. Colored diamonds represent the sampling times analyzed by fluorescence microscopy in the panel B. B) Distribution of GFP fluorescence in the growing population of LF82; GFP fluorescence is expressed as the number of generations (each generation corresponds to a 2-fold decrease in GFP fluorescence compared with the average fluorescence of the fully induced population at t0). C) Distribution of GFP fluorescence in the population of LF82-infecting macrophages. Fluorescence was measured for individual bacteria or small bacterial clusters after macrophage fixation at 1 h, 6 h, 24 h and 48 h post-infection. D) Distribution of the GFP fluorescence in the population of K12-C600 bacteria infecting macrophages. Fluorescence was measured for individual bacteria or small bacterial clusters after macrophage fixation at 1 h and 24 h post-infection.

**Supplementary Figure S5. Calibration of the TIMER assay.** A) Scatter plot of green versus red TIMER fluorescence measured for exponentially growing LF82 (green) and for a culture that had reached stationary phase (red). B) Distribution of the red TIMER fluorescence (Arbitrary fluorescence Intensity, AI) measured for exponentially growing LF82 (green) and stationary phase LF82 (red) at 4 h and 18 h post-infection. Curves represent the normal fit of the data. The middle peak height for exponential and stationary cultures was used to respectively define the fast-mid and mid-slow borders of the boxes.

