## supplementary informations for "The Crohn’s disease-associated *Escherichia coli* strain LF82 rely on SOS and stringent responses to survive, multiply and tolerate antibiotics within macrophages"

**Legend of the supplementary figures**

**Supplementary Figure S1**. A) Growth curves of LF82 and K12 (C600) in LB medium at pH 7.4 (blue) and in the presence of SHX (15 mg/ml), SHX (7 mg/ml), EDTA (70 mM) or in LB at pH 4.7 and LB at pH 4.7 in the presence of EDTA (70 mM). B) Chemicals or a pH shift were applied at 160 min. B) same as in A with addition of the antibiotics ciprofloxacin (24 ng/ml) or cefotaxim (800 ng/ml). Data are the mean of 3 technical replicates.

**Supplementary Figure S2.** A) Live and dead assay performed in situ on infected macrophages and after macrophage lysis. For in situ experiments a very weak propidium iodide (PI) labeling is observed on putative dead LF82. By contrast strong PI labeling is observed after macrophage lysis. B) Measure of the speed of disappearance after phagocytosis by macrophages of heat-killed LF82. LF82 were killed by 15 min incubation at 60°C and subsequently labeled with propidium iodide. Labeled dead LF82 were incubated with macrophage at an MOI of 100. Imaging was performed at 1h, 2h, 3h and 24h post infection. SYTO-9 was used to reveal macrophage and eventual live bacteria. The number of dead LF82 per macrophage was measured at each time points. Data are average of 3 experiments.

**Supplementary Figure S3**. A) Measure of the resistance to MMC of LF82, LF82 *recA*, LF82 *lexAind-,* MG1655, MG1655 *recA* and MG1655 *lexAind-*. B) Measure of the tolerance to cefotaxim of LF82 and LF82 *recA*, LF82 *lexAind-*, LF82 *sulA* grown in LB medium to an OD of 0.2. C) Induction of tolerance to cefotaxim by pretreatment with a subinhibitory dose of ciprofloxacin (24 ng/ml). The data represent the ratio of the number of bacteria that were tolerant to 3 h of cefotaxim in the presence or absence of ciprofloxacin.

**Supplementary Figure S4**. A) Growth curve of LF82 pFC6Gi in LB at 37°C. Colored diamonds represent the sampling times analyzed by fluorescence microscopy in the panel B. B) Distribution of GFP fluorescence in the growing population of LF82; GFP fluorescence is expressed as the number of generations (each generation corresponds to a 2-fold decrease in GFP fluorescence compared with the average fluorescence of the fully induced population at t0). C) Distribution of GFP fluorescence in the population of LF82-infecting macrophages.

Fluorescence was measured for individual bacteria or small bacterial clusters after macrophage fixation at 1 h, 6 h, 24 h and 48 h post-infection. D) Distribution of the GFP fluorescence in the population of K12 bacteria infecting macrophages. Fluorescence was measured for individual bacteria or small bacterial clusters after macrophage fixation at 1 h and 24 h post-infection.

**Supplementary Figure S5**. A) Scatter plot of green versus red TIMER fluorescence measured for exponentially growing LF82 (green) and for a culture that had reached stationary phase (red). B) Distribution of the red TIMER fluorescence measured for exponentially growing LF82 (green) and stationary phase LF82 (red) at 4 h and 18 h post-infection. Curves represent the normal fit of the data. The middle peak height for exponential and stationary cultures was used to respectively define the fast-mid and mid-slow borders of the boxes.

Supplementary Figure S1

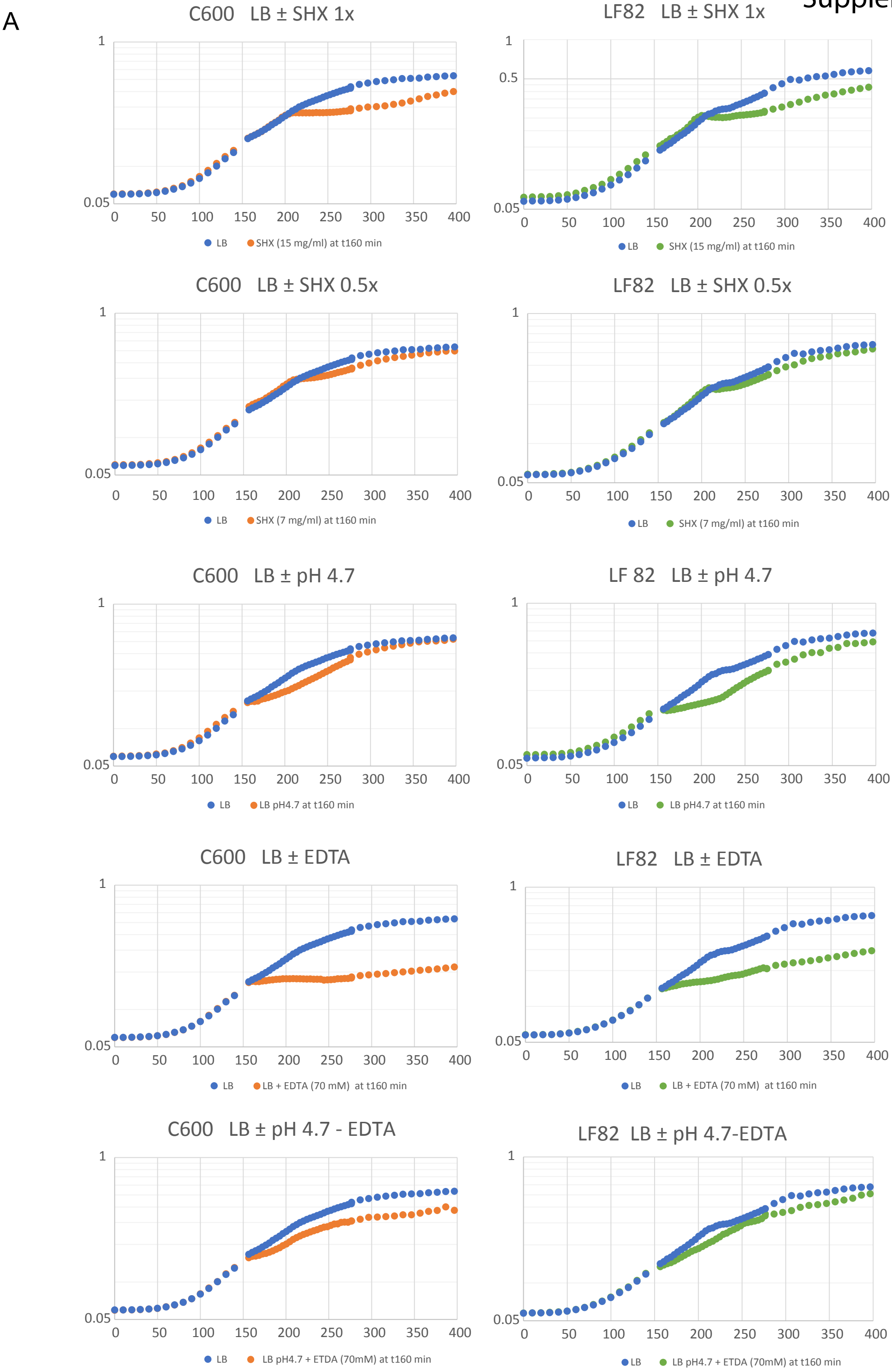

B

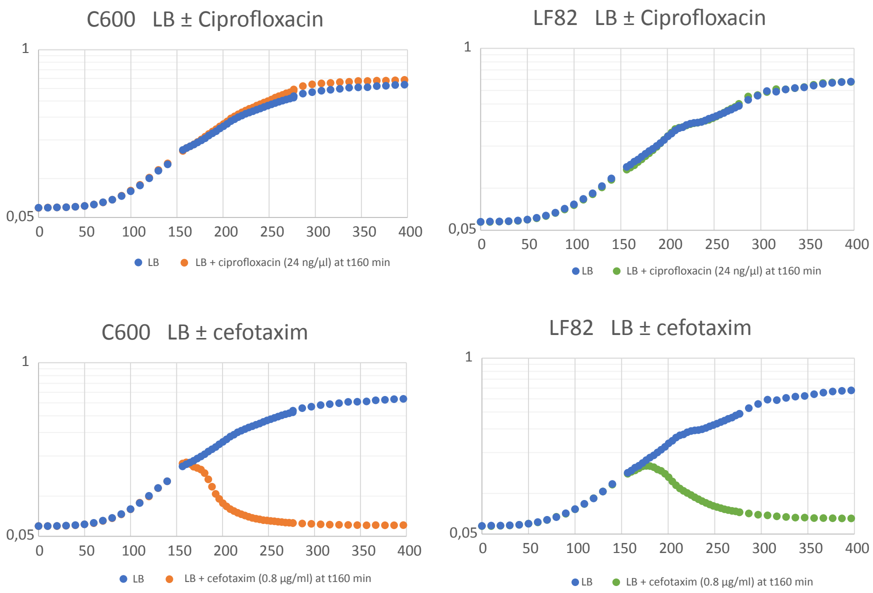
C600 LB ± Ciprofloxacin
1
0,05
0 50 100 150 200 250 300 350 400
LB
LB + ciprofloxacin (24 ng/µl) at t160 min
LF82 LB ± Ciprofloxacin
1
0,05
0 50 100 150 200 250 300 350 400
LB
LB + ciprofloxacin (24 ng/µl) at t160 min
C600 LB ± cefotaxim
1
0,05
0 50 100 150 200 250 300 350 400
LB
LB + cefotaxim (0.8 µg/ml) at t160 min
LF82 LB ± cefotaxim
1
0,05
0 50 100 150 200 250 300 350 400
LB
LB + cefotaxim (0.8 µg/ml) at t160 min

Supplementary Figure S2

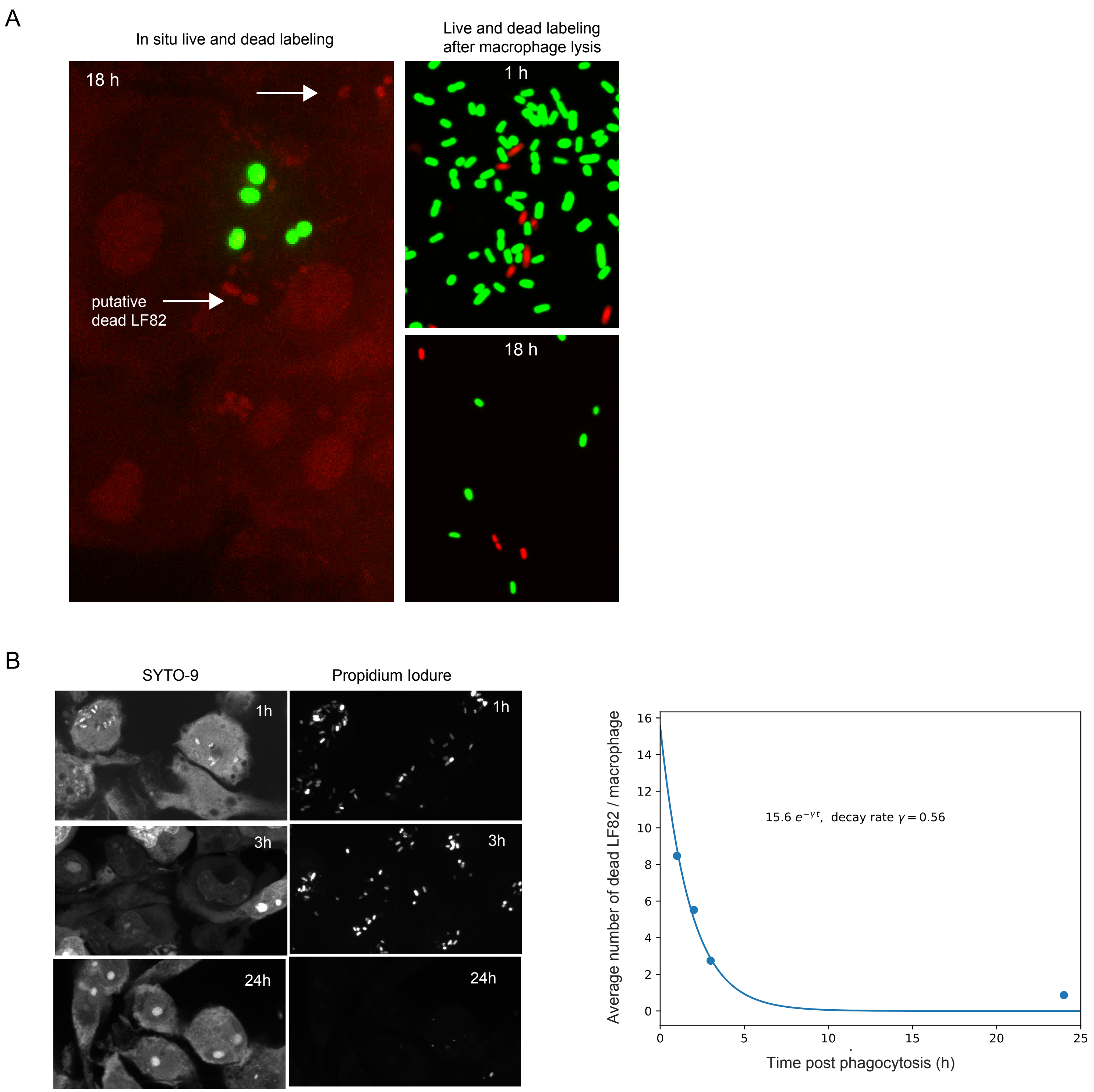

### Supplementary Figure S3

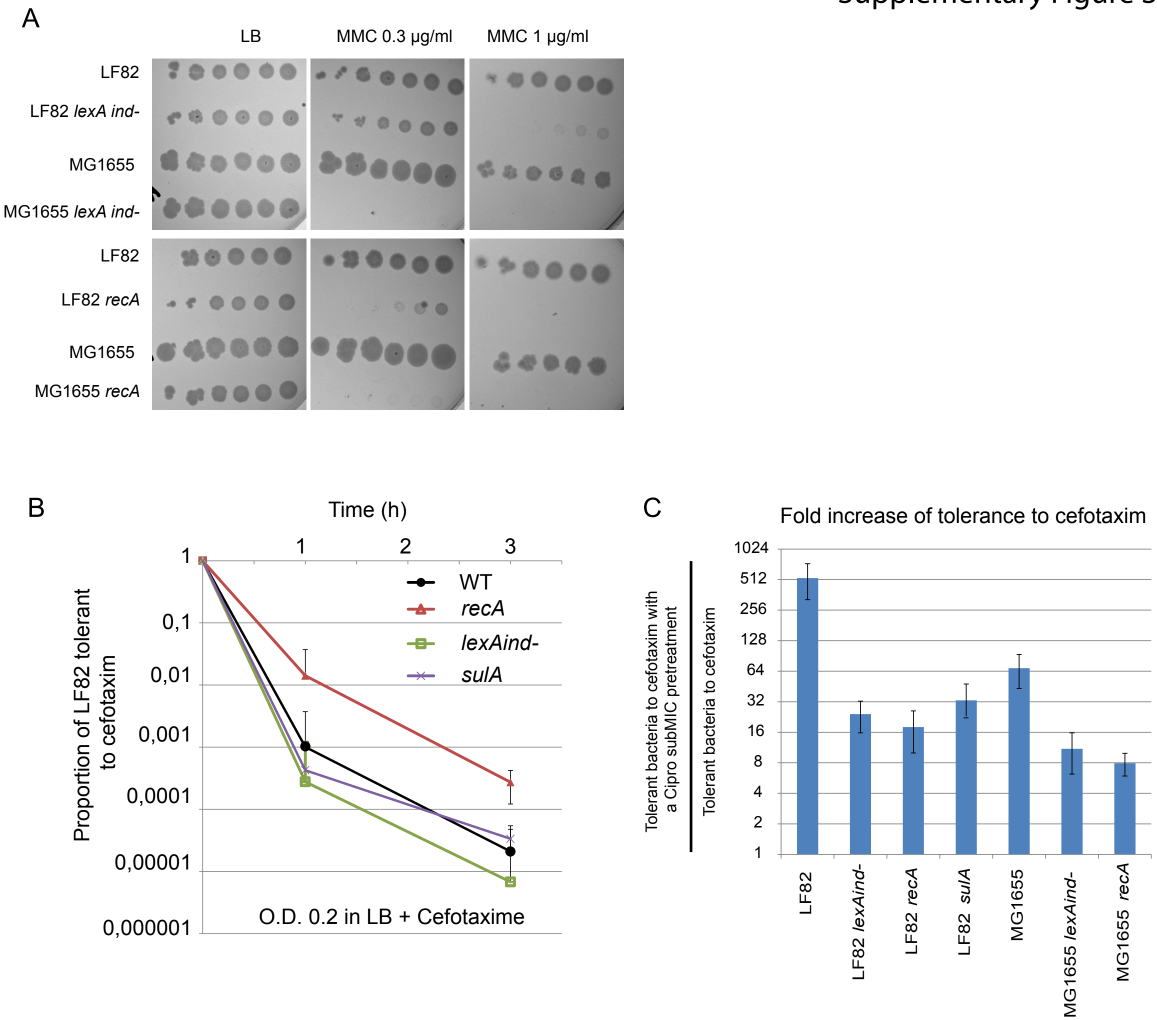

Supplementary Figure S4

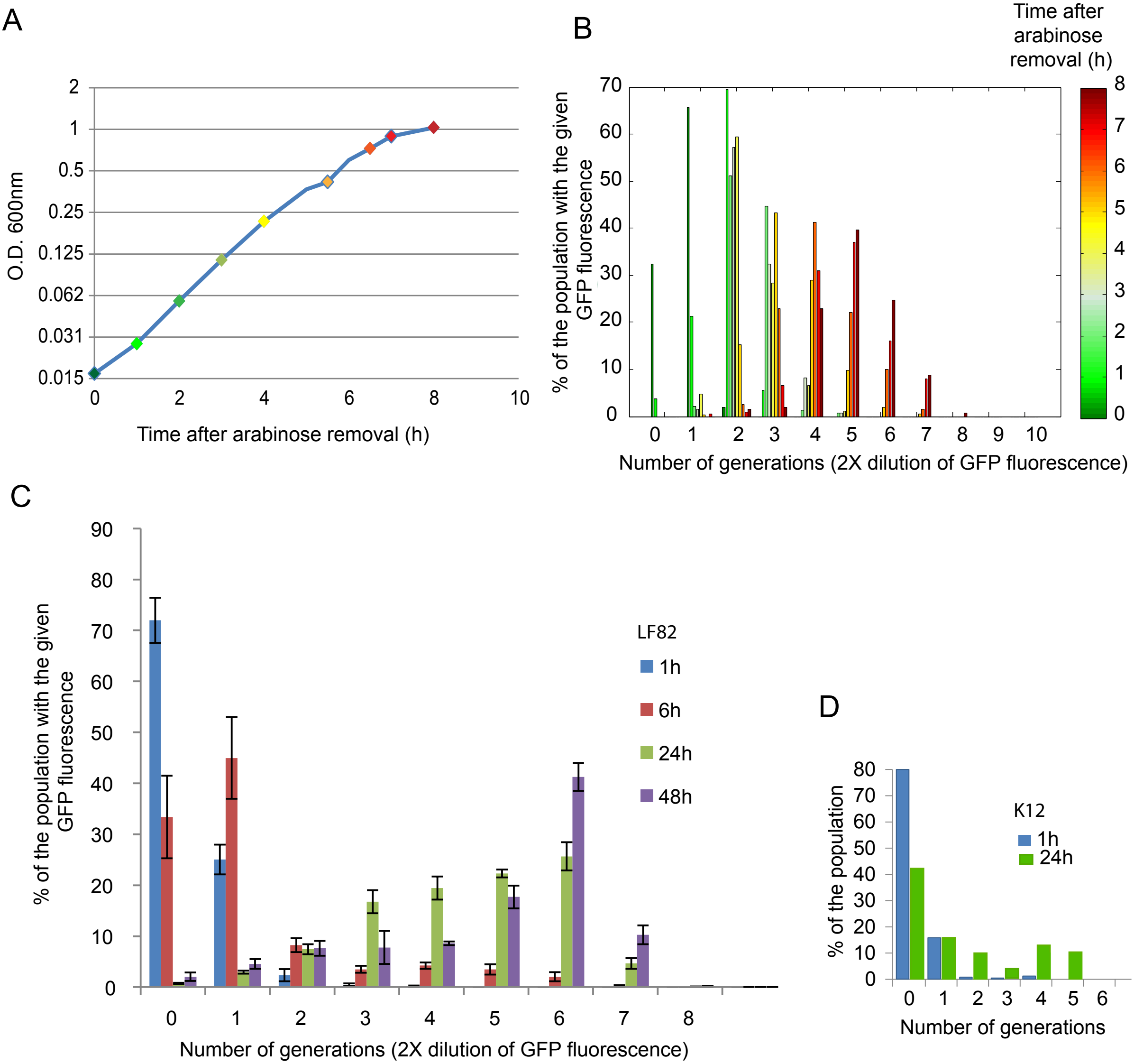

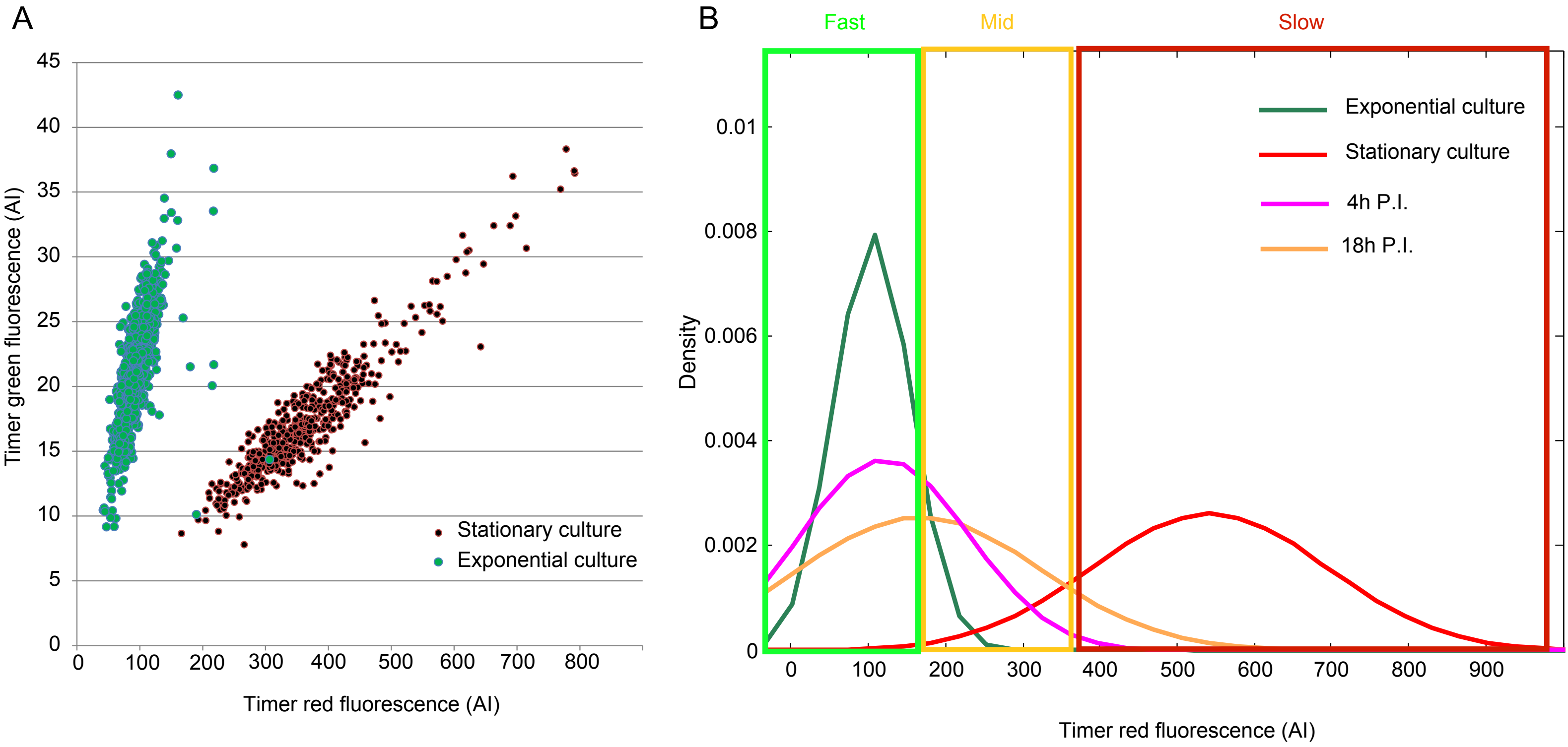
A
Timer green fluorescence (AI)
Timer red fluorescence (AI)
Stationary culture
Exponential culture
B
Fast
Mid
Slow
Density
Exponential culture
Stationary culture
4h P.I.
18h P.I.
Timer red fluorescence (AI)

### Supplementary material S1

**1 The model** **1**

**2 Data** **3**

**3 Fit of the data with the model** **5**

**4 Parameter variation** **9**

### 1 The model

The biological observations on the differential behaviour in macrophage infection by *E. coli* K12, the AIEC strain LF82 and its stringent response mutant LF82*dksA* can be compounded in a mathematical model describing the dynamics of the bacterial population. This model can in turn be used to provide a quantitative estimate of biologically important rates, such as net growth rate within the macrophage and rate of persister generation, for the different strains.

The model has three state variables, that change in time: the number of non-persistent $B$ and persistent $P$ bacteria, and the level of stress $S$. The number of dead bacteria $D$ can be computed based on those variables. The experimental observables are expressed as functions of the three state variables: the total population size of living bacteria is $B+P$, the relative amount of persisters is $P/(B+P)$ and the number of dead bacteria is $D$. The stress level, on the other hand, is not directly measurable but the relevance of stress in this context can be inferred from observations of permanent expression of stress-related genes, together with the effect of their mutations on bacterial proliferation.

The model recapitulates the dynamics of infection within the macrophage and is temporally structured in a lag and a growth phase (illustrated in Figure 6 A). At time t=0, when bacteria from an exponentially growing culture enter the macrophage, the impact with the new challenging environment results in a first wave of immediate death, occurring within 1 h post infection (P.I.) (Figure 1 A). The model ignores the bacteria killed in this early event. Since the fraction of persisters is small in *in vitro* cultures, we assume that surviving cells are all non-persisters, so that $P(0) = 0$. Moreover, we choose $B(0) = 1$ without loss of generality (corresponding to the normalization in Fig. 1 A). This way, comparison in the kinetics for different strains is independent of the initial number of cells that got killed immediately. We also consider that at the beginning of the infection stress is negligible ($S = 0$).

Following entry in the phagolysosomes, we describe the population dynamics as the succession of a lag period where population net growth is nonpositive, followed by an expansive phase where cells actively divide. The existence of a lag phase is widely observed following a sudden environmental shift (here the entry into the macrophage), notably when this poses a significant challenge to survival. In our model, the duration $\lambda$ of the lag phase can differ among strains. During the first hours of experimental observations, K12 and LF82 maintain an almost constant total population size, while LF82*dksA* slightly declines (Fig. 6 B). In lag phase, thus, the net growth rate is $\delta_1 = 0$ for the former two strains and amounts to a death rate $\delta_1 = 0.16$ h$^{-1}$ for the latter. This net death rate is estimated from the additional proportion of death cells in the stringent response mutant with respect to the LF82 strain (Fig. 5 F): supposing that death at infection is the same (15%) for LF82 and its mutant, the latter undergoes an additional 15% death in the first hour, thus $\delta_1 = -\ln(1 - 0.15) = 0.16$.

At the exit of the lag phase, the population resumes a positive net growth rate (the difference of an unknown, possibly larger, gross birth rate and an unknown gross death rate), whose maximum value $\beta$ is attained in exponential phase. If we single out the exponential phases of the growth curves, we find that the maximal net growth rate is comparable for the three strains (although we only disposed of two points for K12), so we assume that $\beta$ is strain-independent.

We consider that stress builds up all along the course of the infection, for instance through accumulation of DNA damage, toxification of the phagolysosome environment, or response of macrophages to the infection. The prolonged permanence within crowded phagolysosomes generates long-lasting effects that can decrease division rate, increase death rate, or both. We model such stress via a dimensionless variable $S$ that increases proportionally to the total bacterial population size, and effectively integrates the number of bacteria that participate to the infection over time. Stress induces death (or reduces division) at a rate $\delta_2(S)$. Negligible in the first phases of the infection, stress-induced death becomes dominant after exponential population growth, and can lead to the downturn of population size in the last few hours of the experiment. A death rate linearly increasing with stress was insufficient to quantitatively reproduce the population decrease, so we introduce a sigmoidal form for the stress-induced death rate:

$$\delta_2(S) = \frac{d_{max}}{1 + e^{a(S_{1/2} - S)}}, \tag{1}$$

where $d_{max}$ is the strain-dependent maximum stress-induced death rate, $S_{1/2}$ measures the stress intensity for which death rate is half its maximal value, and $a$ is the sensitivity of the nonlinear response to a change in stress level (higher levels of $a$ produce a response that is more step-like). For simplicity, we assume that the last two parameters are the same in all strains, so that the differences in stress response are chiefly determined by the maximal death rate.

According to the previously stated assumptions, the dynamics of a well-mixed population of bacteria can be described as the solution of the following system of ordinary differential equations:

$$\begin{aligned} \frac{dB}{dt} &= [1 - \mathbb{1}_{t<\lambda}]\, \beta\, B - \mathbb{1}_{t<\lambda}\, \delta_1\, B - \delta_2(S)\, B - k_p\, B \\ \frac{dP}{dt} &= k_p\, B \\ \frac{dS}{dt} &= B + P. \end{aligned} \tag{2}$$

The indicator function $\mathbb{1}_{t<\lambda}$ is one during the lag phase (then the first term in $\frac{dB}{dt}$ cancels) and zero otherwise (then the second term cancels), and provides a compact representation of the existence of a lag phase (for $t < \lambda$) and an expanding phase (for $t > \lambda$).

We can moreover compute the number $D$ of dead cells by solving a fourth, subordinate, equation:

$$\frac{dD}{dt} = \mathbb{1}_{t<\lambda}\, \delta_1\, B + \delta_2(S)\, B - \gamma\, D, \tag{3}$$

where the clearance rate of dead cells has been estimated to $\gamma = 0.6$ h$^{-1}$ by fitting the decay of heat-killed bacteria in macrophages (Fig. 3 B).

For each strain, the infection dynamics is obtained as a solution of eqs. 2, which depend on 6 parameters for a single strain, or 12 parameters if all strains are simultanaously considered.

The model is implemented in the python programming language, and numerical integration realised through `odeint` (from the `scipy integrate` module) with a maximum integration step size of $\Delta t = 0.1$.

#### 2 Data

We dispose of measurements of colony forming units (CFU), that are obtained after bacteria have spent a time $t$ in the macrophage (see Methods in the main text, 3 to 12 replicates depending on the time point). The number of CFU approximates the total number of bacteria $B(t) + P(t)$. Figure S1.1 displays the individual data points (normalized so that the mean initial population size is 1 for all strains), their mean and standard deviation.

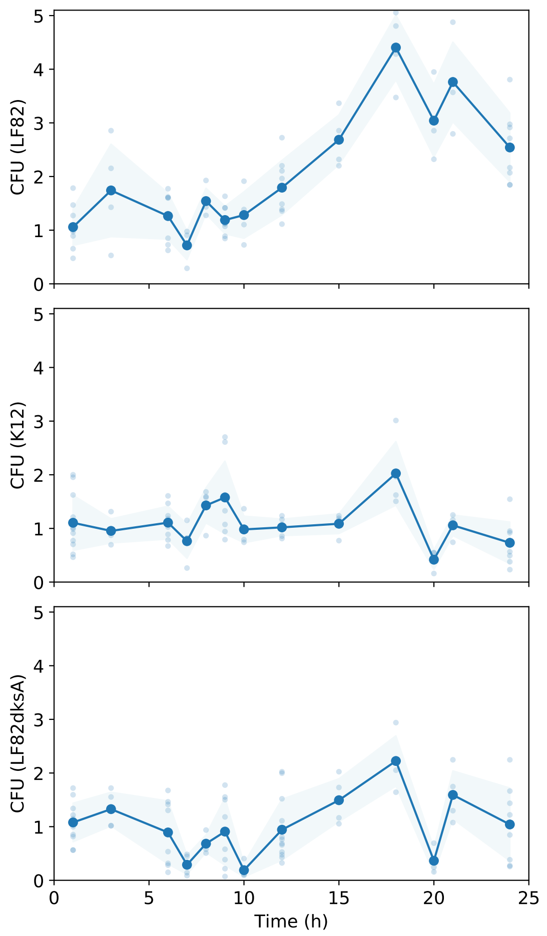


Figure S1.1: **Total normalised number of bacteria in macrophage in the course of the infection.** The infection kinetics is quantified by observing CFUs for the three strains *E. coli* LF82, K12 and LF82*dksA*. Small dots represent individual experiments. Their means are shown as a large dots and standard deviations as a shaded area. These statistics are also reported in Fig. 6B

Moreover, we have measures of relative abundance of bacteria extracted from macrophages at 1, 6, 20 and 24 h PI that are tolerant to 3 h exposure to the antibiotic ciprofloxacin (Fig. 4 A and Methods), recapitulated in Table S1.1. From these frequencies and the total population size data, we can compute the number of persisters. Since persister quantification involves a considerably larger experimental effort, these data have a lower temporal resolution than CFU counts.

| $t$ (h) | 1 | 6 | 20 | 24 |
|---|---|---|---|---|
| LF82 | 0.0262 | 0.0433 | 0.0963 | 0.1359 |
| K12 | 0.0022 | 0.0154 | 0.0126 | 0.0565 |
| LF82kdsA | 0.0052 | 0.0033 | 0.0268 | 0.0544 |

Table S1.1: **Relative abundance of persisters.** Relative amount of the bacteria extracted from macrophages $t$ hours PI, that are able to form colonies after 3 hours exposure to antibiotic treatment.

In the fitting procedure, we weigh the number of persisters so that it is commensurable with the population size (which is itself rescaled by normalization). Since the number of persisters is a small percentage of the total population size, the mean square distance of the data from the simulated trajectory is one order of magnitude smaller for the persister measures than for the total population size data. Combined with the lower temporal resolution of the former, the fit would be biased towards the optimization of the population size data points. We have solved this problem by a normalization that brings population size and persister number on the same scale. Before optimization, we multiply the number of persisters for all strains with the ratio between the mean population size and the mean number of persisters measured for all strains (factor 19.2).

For the pathogenic LF82, we further know the relative amount of dead cells present in macrophages at several time points P.I.. Since the total number of bacteria $B + P$, as measured by CFU counts, only includes viable bacteria, the relative amount of dead cells is $\kappa = \frac{D}{B+P+D}$. With this assumption we can estimate the total number of dead cells comparable with CFU counts $D = \frac{\kappa}{1-\kappa}(B + P)$. This data is subjected to multiple sources of error, and the model estimates are themselves dependent on the use of a decay rate constantly equal to that observed for heat-killed cells. Therefore, we do not include these data in the model fit, but use them for a qualitative comparison with the model predictions.

| $t$ (h) | 1 | 10 | 18 | 24 |
|---|---|---|---|---|
| mean $\kappa$ | 0.21 | 0.19 | 0.28 | 0.32 |
| std dev | 0.047 | 0.041 | 0.028 | 0.248 |

Table S1.2: **Relative abundance of dead LF82.** Relative amount of dead bacteria $t$ hours PI and standard deviation of those measures.

### 3 Fit of the data with the model

Since different processes described by the model are important for different phases of the infection dynamics, we have devised a two-step fitting method that allows us to assess first the parameters involved in lag and exponential phase of growth, and then those relative to stress-induced death. The latter are only relevant for the last few data points, hence their estimation is less precise than for the other parameters, and can only be used for qualitative, and not a quantitative, comparison among strains.

During the first 18 hours post-infection, when stress-induced death is negligible ($\delta_2(S) = 0$), eqs. 2 can be analytically solved to yield the dynamics of the total population size and of the persister component:

$$
\begin{aligned}
B(t) &= \begin{cases} B_0\, e^{-(\delta_1+k_p)t} & \text{for } t < \lambda \\ B_\lambda\, e^{(\beta-k_p)(t-\lambda)} & \text{else} \end{cases} \\
P(t) &= \begin{cases} B_0\, \frac{k_p}{\delta_1+k_p}\, \left[1 - e^{-(\delta_1+k_p)t}\right] & \text{for } t < \lambda \\ P_\lambda + B_\lambda\, \frac{k_p}{k_p-\beta}\, \left[1 - e^{(\beta-k_p)(t-\lambda)}\right] & \text{else} \end{cases}
\end{aligned}
\tag{4}
$$

with $B_\lambda = B(\lambda) = B_0\, e^{-(\delta_1+k_p)\lambda}$ and $P_\lambda = P(\lambda) = B_0\, \frac{k_p}{\delta_1+k_p}\, \left[1 - e^{-(\delta_1+k_p)\lambda}\right]$.

These equations depend on seven parameters: the growth rate $\beta$, common to all strains, and the strain-specific lag phase duration $\lambda$ and rate of persister production $k_p$. In the first step of the fit, we estimate these parameters by minimizing the quadratic distance between eq. 4 evaluated at the times of observation and the data, after persister numbers are rescaled as described above. The fit is optimized with the algorithm `least_squares` (`scipy optimize` module) and initial estimates $k_p = 10^{-2}, 10^{-4}, 10^{-3}$, $\lambda = 7, 7, 7$ (for LF82, K12 and LF82*dksA* respectively) and $\beta = 0.2$, with $\lambda$ and $\beta$ bound to positive values and $k_p \in [10^{-10}, 10^{10}]$. We approximate the missing persister measure at 18 h, which was not performed, with that at 20 hours. We consider, that persisters are protected from death on the time scale of the infection, and that their production rate is small, so that their number is not expected to change much in 2 hours. The best fit of this first step is shown in Figure S1.2 A.

In a second step, we fix the parameters obtained in the first step, then use the complete data set and the full model equation (2) to estimate the remaining parameters for stress-dependent death: $S_{1/2}$, $a$ (common to all strains) and the strain-dependent maximal death rates $d_{max}$. The switch rate $k_p$ was also refined with the full data set. The fit is optimized with the `differential evolution` algorithm (`scipy optimize` module with `popsize`= 100, `mutation`= $(0.7, 1)$ and `recombination`= 0.3) with parameters bound to positive values until 40, but with $k_p$ between $10^{-10}$ and $10^{10}$ for faster convergence. The best fit in this second step is illustrated in Figure S1.2 B.

We then evaluated the uncertainty in the parameter estimates by performing the fit (with `popsize`= 20, `mutation`= $(0.5, 1)$ and `recombination`= 0.5) on one hundred reduced data sets, where one data point (small dots in Figure S1.1) was removed from each set of CFU counts. These synthetic datasets had different mean CFU counts, thus different best fitting curves. The standard deviation of the best fitting parameters provides a measure of the robustness of the estimates with respect to missing observations: it is higher when the quantification critically depends on one of the data points, and is thus considered less reliable. The robustness of $k_p$ switching rates is not representative, since only the total number of CFU, not the amount of persisters, was changed.

Parameters corresponding to the best fitting trajectory and their standard deviation are collected in Table S1.3.

As expected, some parameters are robust, indicating that their quantification is based on sufficient data, while others would require more data for a sound quantitative estimate. The growth rate $\beta$ and the lag times $\lambda$ are the most accurately estimated.

All strains grow at a much slower net rate during the infection with respect to growth in lysogeny broth (LB). This is expected, as the macrophage environment is likely to be not only nutrient-poor, but also filled with stressors. The estimated **maximum net growth rate** $\beta = 0.15$ h$^{-1}$ corresponds to the

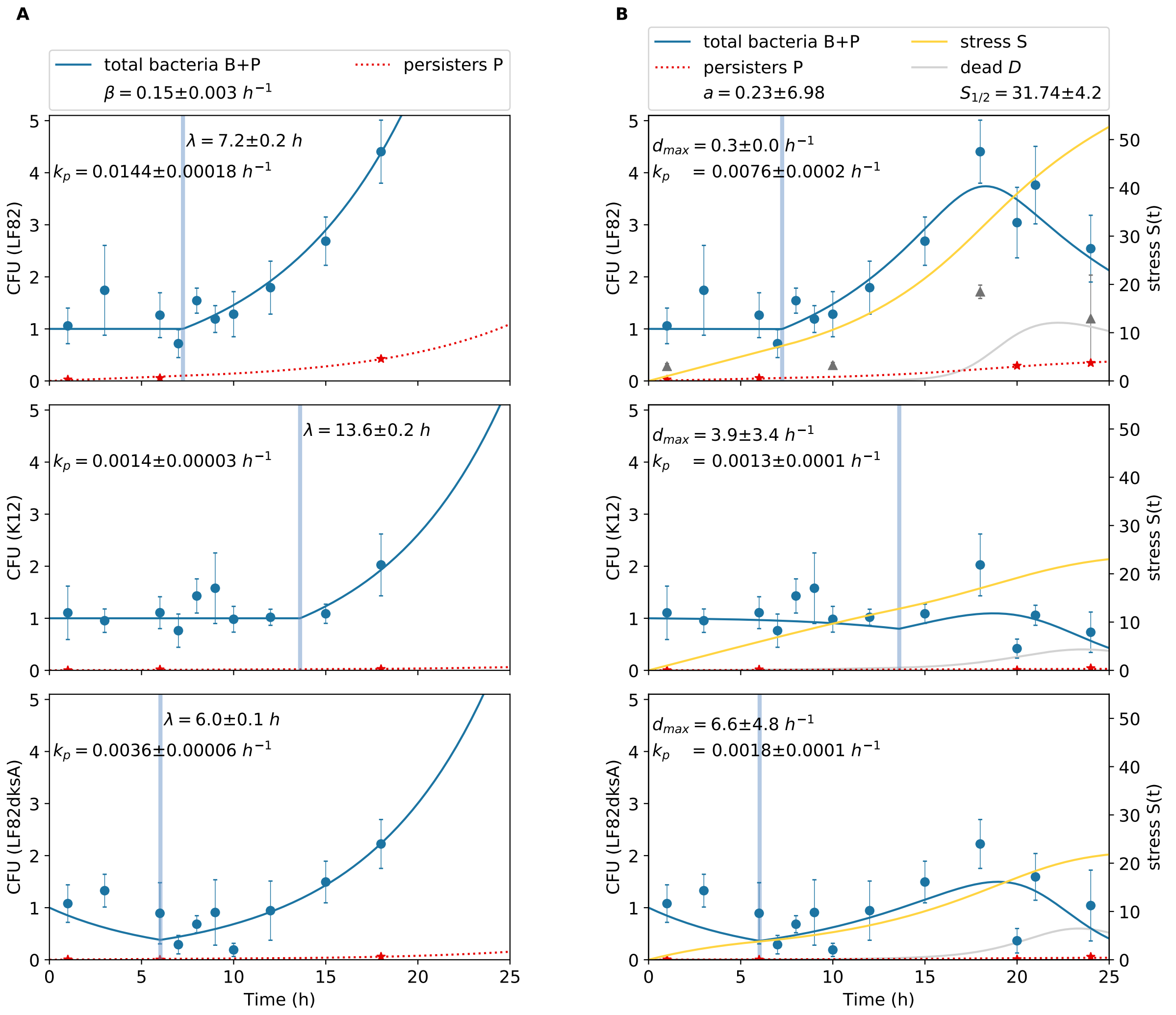


Figure S1.2: **Two-steps data fit**. (**A**) Step 1 of the fitting method explained in the text embraces lag phase and exponential growth up to $t = 18$ h. Experimental measures of CFU and persister counts are fitted with equations (4), obtained from equations (2) when stress-dependent death rate is negligible ($\delta_2(S) = 0$). The best fit provides estimates for the net growth rate $\beta$ (assumed to be the same for all strains), the lag phase duration $\lambda$ and the rate of switch to persisters $k_p$. During the lag period only LF82*dksA* experiences a non-zero net death rate $\delta_1 = 0.16$ h$^{-1}$. (**B**) Step 2 of the fit allows to estimate the parameters defining the stress dependent death rate $\delta_2(S)$: $d_{max}$, $a$ and $S_{1/2}$. While the parameters obtained in step 1 are held constant, stress-related parameters and again $k_p$ are estimated by fitting data points up to 24 hours with the eq. (2). The number of dead cells, as defined in Eq. (3), is compared with the approximate normalised number of dead cells, calculated from the percentage of dead cells (see Table S1.2 and main text Fig 1A). Table S1.3 compiles the parameters corresponding to the best fit.

division rate minus a possible constant mortality rate in the phagolysosome environment. It corresponds to a doubling period $T_{double} = \log 2/\beta = 4.6$ h, which is comparable to the estimations obtained by

| Bacterium | $\lambda$(h) | $\beta$ ($h^{-1}$) | $\delta_1$ ($h^{-1}$) | $k_p$ ($h^{-1}$) | $a$ | $S_{1/2}$ | $d_{max}$ ($h^{-1}$) |
|---|---|---|---|---|---|---|---|
| LF82 | 7.2±0.2 | | *0* | 0.0076±0.0002 | | | 0.3±0.0 |
| K12 | 13.6±0.2 | 0.15±0.003 | *0* | 0.0013±0.0001 | 0.23±6.98 | 31.74±4.2 | 3.9±3.4 |
| LF82*dksA* | 6.0±0.1 | | *0.16* | 0.0018±0.0001 | | | 6.6±4.8 |

Table S1.3: **Parameters from the fit.** The death rate in lag phase, in italics, is set to 0 for K12 and LF82, while it is $\delta_1 = 0.16$ for the mutant LF82*dksA*. The other parameters correspond to the fit obtained through the two-step method explained in the text. Times smaller than 18 hours PI allow us to fit: the net growth rate $\beta$, common for all strains, the lag time $\lambda$ and switch rate $k_p$ for each strain. Parameters for the non-linear death rate $\delta_2(S)$, with sensitivity $a$ and $S_{1/2}$ equal for all strains, but different maximum death rates $d_{max}$, are subsequently fitted including all time points. The final value of $k_p$ is also obtained with the full data set.

fluorescent dilution (Supp. Fig. 2), and more than five times smaller than the maximum net growth rate measured for LF82 in LB (doubling period 50 minutes).

The fit evidences a big difference in **lag phase duration** $\lambda$ among strains, and notably between the commensal K12 strain on the one hand, and the pathogen LF82 and its mutant on the other. Pathogens have a head start in the infection, as they exit lag phase in about half the 13 hours necessary for K12. Such reduction in lag duration does not appear to be lost in the mutant, in spite of stringent response being impaired. The precision of our estimates however does not allow to firmly conclude about possible variations in lag duration between the two LF82 strains. Small differences between their estimates might moreover be affected by the normalization of the CFU counts to the measure at $t = 1$ h, that does not correct for the observation error and may cause an underestimation of the population size in lag phase for the LF82 strain. Furthermore, a differential estimation of growth rates, which we assumed to be equal here (but this is based on circa 2 data points for K12), could also change the length of lag period between K12 and LF82.

Even though the lower number of data points makes their estimate more uncertain, the fit reveals important differences in the **rate of switch to persistence** $k_p$ of the different strains, consistent with the differences observed in vitro. The pathogenic LF82 strain has a rate of persister production $k_p = 0.0076$, six times larger then the commensal K12 strain, and the stringent response mutant produces persisters at a rate slightly higher than K12, but still much lower rate than LF82, suggesting a possible involvement of the stringent response in the capacity of developing antibiotic tolerance.

Based on the the rate of switch and the doubling time, we can compute an upper estimate for the likelihood that a bacterium becomes a persister during its lifetime. If $\beta$ measures the actual net rate of birth, the probability of never switching to a persister is $p_{no\,switch} = (1-k_p)^{T_{double}}$. Hence, the probability of switching is $p_{switch} = 1 - p_{no\,switch}$, that is 3.5 % for LF82, 0.6 % for K12 and 0.8 % for LF82*dksA*.

The **stress-dependent death rate**, corresponding to the estimated values of $d_{max}$, $a$ and $S_{1/2}$, is visualised in Figure S1.3 A. The time evolution of the experienced stress and the resulting death rate are shown in Figure S1.3 B.

The three strains differ in their maximum death rate and thus their susceptibility to stress. Because of the uncertainty of the estimate (few data points, larger errors), the strong assumption that stress is integrated over bacterial abundance, and the fact that the stress levels in K12 and LF82*dksA* remain relatively small, however, it is impossible to go beyond a qualitative conclusion: while K12 and the mutant LF82*dksA* appear to be more sensitive to stress, LF82 seems to be able to tolerate much higher values of $S$.

From the fit above we can derive the amount of dead cells present during the infection kinetics (eq. (3)). For the wildtype strain LF82 we have measured the relative amount of dead cells (Table S1.2) present in the macrophage for some time points. From this we calculate the absolute normalised number and compare it to our model result for $D$. Although a quantitative prediction is not possible, the curve matches qualitatively, notably in the prediction of a fast increase in dead cells delayed with respect to the

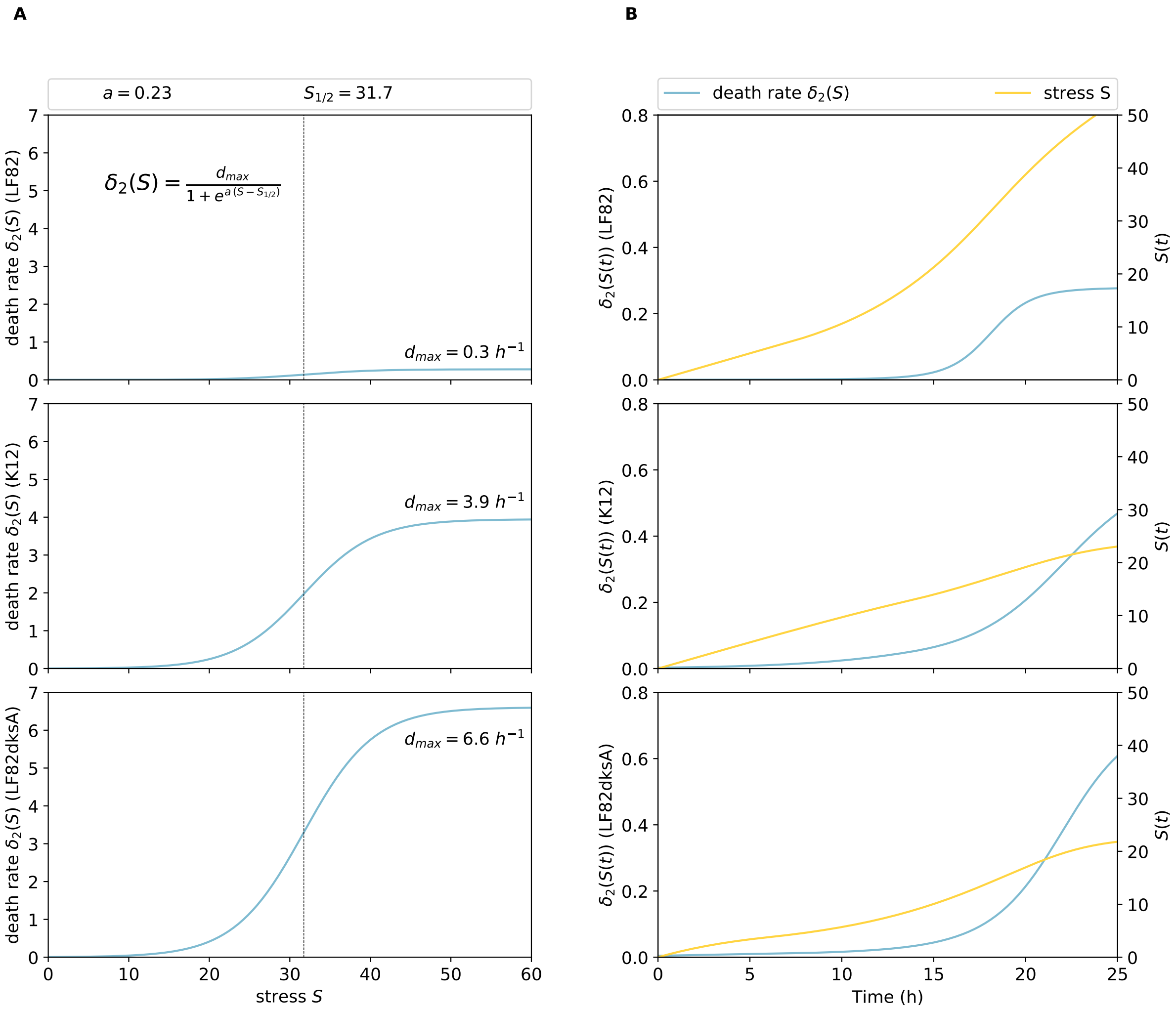


Figure S1.3: **Stress-dependent death rate from fitted parameters.** $\delta_2(S)$ is computed from the parameters $d_{max}$, $a$ and $S_{1/2}$ fitted in step 2. The death rate is displayed as a function of stress $S$ (**A**) and as a function of time $t$ (**B**) along a simulation trajectory. Stress increases progressively, as illustrated in Figure S1.2.

total population size, and a successive decline. One possible explanation for the quantitative discrepancy is the neglect of a component of death that we cannot estimate: that hidden in the net growth rates. Indeed, the net turnover rate is actually the sum of a larger doubling rate and a stress-independent death rate. By considering only cells that are killed by stress, the model thus systematically underestimates the number of dead cells. A second reason for the discrepancy may be a wrong estimation of the value of the decay rate $\gamma$ for dead cells. This was derived from the decay of heat-killed cells measured in uninfected macrophages, and might be lower, or even time-dependent, in macrophages that are under attack.

### 4 Parameter variation

We have seen that differences in lag phase duration and rate of persister generation accounts for most of the discrepancy observed in the infection kinetics between the commensal K12 strain and the pathogenic LF82 strain. We can now use the model to inquire what is more generally the impact of parameter variation onto the infection kinetics. We have therefore produced different trajectories changing one parameter at a time in a range including the values previously fitted.

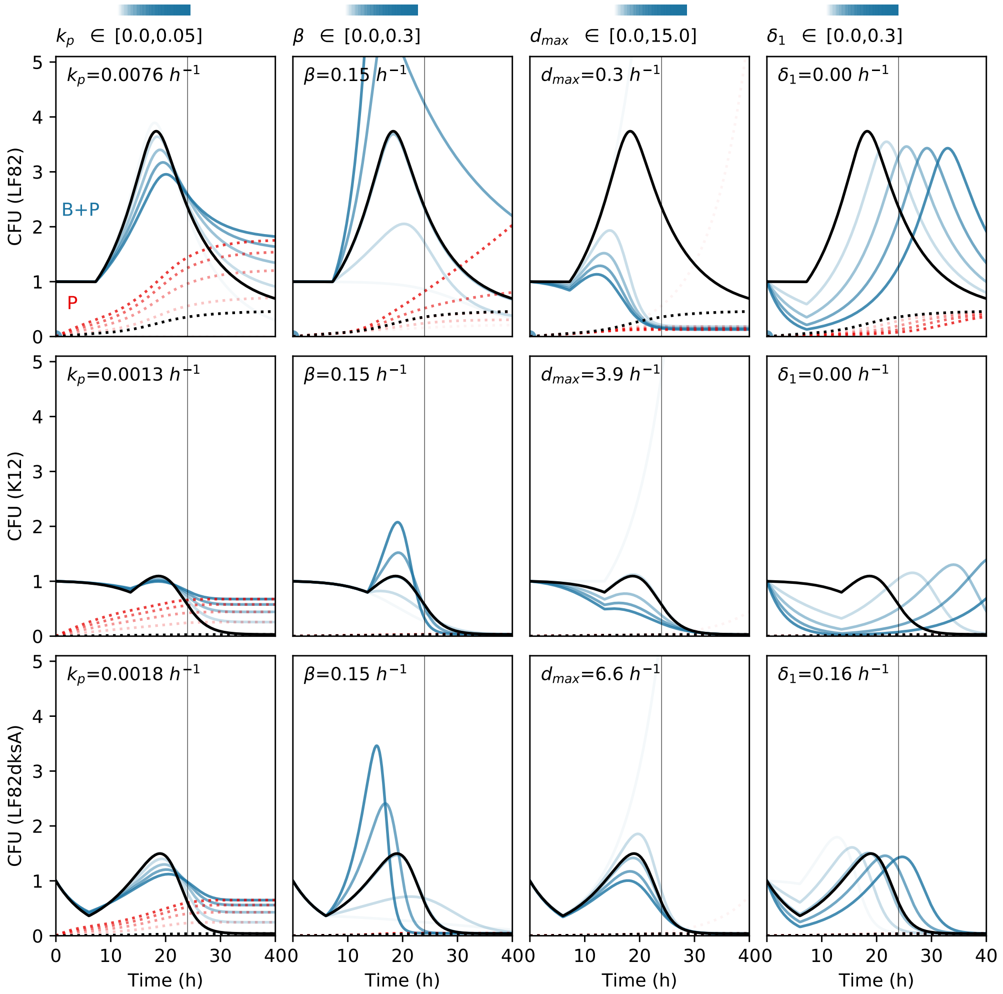


Figure S1.4: **Trajectories of the model when parameters are varied.** Numerical integration of the full model eqs. 2 changing one of the parameters $k_p$, $\beta$, $d_{max}$ or $\delta_1$ in the intervals indicated on top of each column. The remaining parameters are fixed as in Table S1.3. Different rows correspond to different strains: first row, LF82; second row, K12; third row, LF82*dksA*. The curve corresponding to the best fit is plotted in black. In the trajectories simulated with parameter variations, total abundance is represented by the blue lines and persister abundance by the red dotted lines. The opacity of the colour reflects the value of the changing parameter as shown at the top of each column. The vertical grey line indicates 24 hours post-infection, the reference value when observations of Figure S1.5 are realized.

Figure S1.4 shows the trajectories of the total number of bacteria $B(t) + P(t)$ and the number of persisters $P(t)$ corresponding to the variation of one parameter in the range displayed above each column. The reference value for the parameter – relative to the best fit of LF82 kinetics (black line) – is indicated in the corresponding panel. For a given line (strain) these reference values correspond to those listed for the corresponding strain in Table S1.3.

Alternatively, we can fix a time of observation and look at the change in the value of the observables when a parameter is changed as previously described. Figure S1.5, for instance, displays the value of total population size and number of persisters 24 hours post-infection.

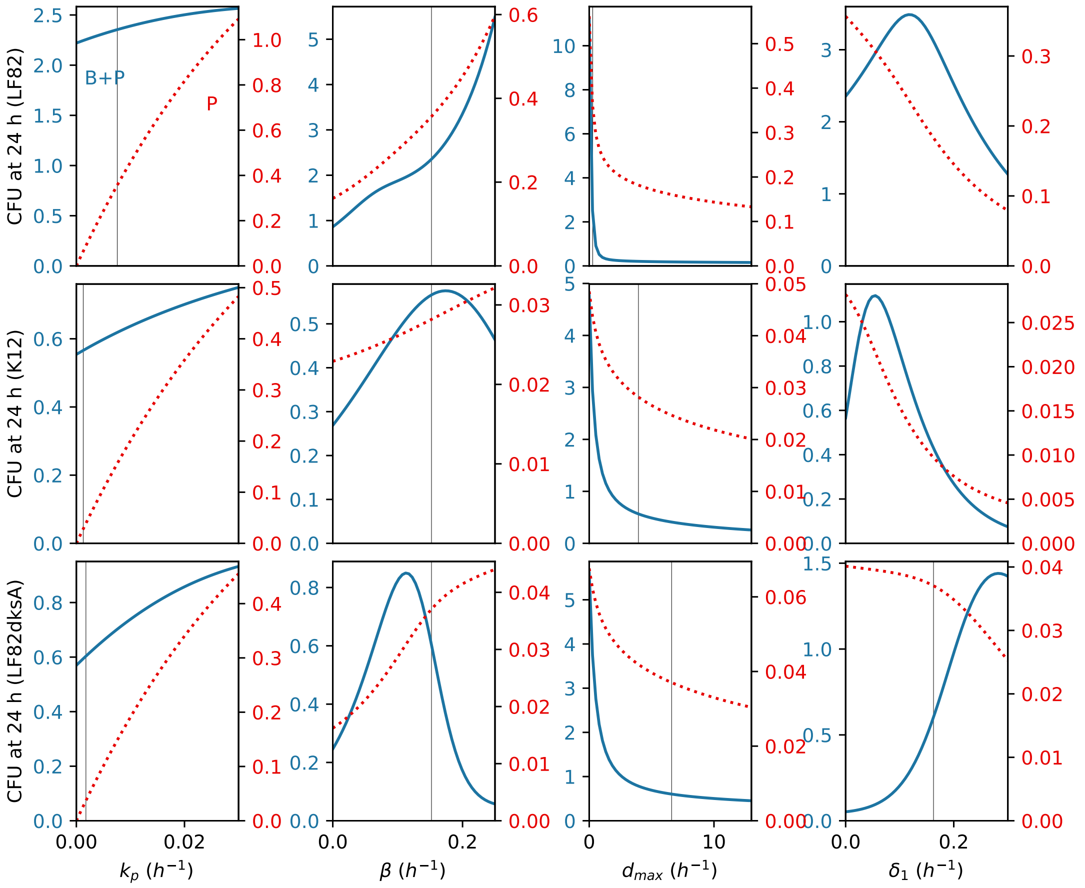


Figure S1.5: **CFU after 24 h from simulations when parameters are varied.** Similar to Figure S1.4, parameters are changed one at a time, starting from the reference values of Table S1.3 (dotted lines). The number at 24 h PI of the total bacteria (blue, left axis) and the persisters (red, right axis) are plotted as a function of the parameters, as these vary in the same intervals indicated in Figure S1.4.

The study of the parameter sensitivity allows us to examine the role of individual parameters in shaping the infection kinetics, and to predict the impact of their variation on experimentally observable quantities.

A higher **rate of switch to persistence** $k_p$ causes, as one would expect, an increase in the number of persisters $P$ troughout the dynamics. The consequent increase in the non-growing fraction of the population, however, also dampens the population overshoot. A secondary effect of such reduced proliferation is that stress becomes relevant at later times, thus additionally boosting persistence through the ongoing generation of persisters by replicating bacteria. A large number of persisters during later times also reduces the impact of stress-dependent death since persisters are assumed not to be killed by stress. Overall, an increase in persister production thus leads, consistently across strains, to larger total population sizes at 24 hours PI.

Increasing the **net growth rate** $\beta$ enhances the overshoot, and as a consequence of increased ploriferation, the number of persisters. However, it can have contrasting effects on the long-term outcome of the infection. A fast population expansion induces a sudden increase of stress, and the subsequent population crash can cause the long-term persister production to be smaller than expected. An increase in the maximum net growth rate can have non-monotonic effects on the total population (see for instance the cases of K12 and LF82*dksA* in Figure S1.5), thus limiting the possible mid-term advantage that can be gained by faster growth. The effect on the long-term dynamics is always positive, since the only bacteria that survive are persisters and their production is increased with faster growth.

An increase in the **maximum death rate** $d_{max}$ unsurprisingly decreases the total abundance of bacteria. Its impact on the number of persisters is less pronounced for intermediate rates, since many persisters are produced before stress-induced death becomes relevant.

Finally, an increase in the **lag phase death rate** $\delta_1$, which was assumed to be zero for LF82 and K12, delays the dynamics without changing dramatically the peak population size. Since proliferation is pushed towards times when stress had more time to acumulate, the delay results in a small decline in the number of persisters generated in the long term. Depending on which side of the overshoot the observation time point is, however, the total population size can either increase (if the observation occurs after the peak has been reached) or decrease, resulting sometimes in non-monotonic variations (e.g. for LF82 and K12).
